# Universal amyloidogenicity of patient-derived immunoglobulin light chains

**DOI:** 10.1101/2021.05.12.443858

**Authors:** Rebecca Sternke-Hoffmann, Thomas Pauly, Rasmus K. Norrild, Jan Hansen, Mathieu Dupré, Florian Tucholski, Magalie Duchateau, Martial Rey, Sabine Metzger, Amelie Boquoi, Florian Platten, Stefan U. Egelhaaf, Julia Chamot-Rooke, Roland Fenk, Luitgard Nagel-Steger, Rainer Haas, Alexander K. Buell

## Abstract

The deposition of immunoglobulin light chains (IgLCs) in the form of amorphous aggregates or amyloid fibrils in different tissues of patients can lead to severe and potentially fatal organ damage, requiring transplantation in some cases. There has been great interest in recent years to elucidate the origin of the very different *in vivo* solubilities of IgLCs, as well as the molecular determinants that drive either the formation of ordered amyloid fibrils or disordered amorphous aggregates. It is commonly thought that the reason of this differential aggregation behaviour is to be found in the amino acid sequences of the respective IgLCs, i.e. that some sequences display higher intrinsic tendencies to form amyloid fibrils. Here we perform in depth Thermodynamic and Aggregation Fingerprinting (ThAgg-Fip) of 9 multiple myeloma patient-derived IgLCs, the amino acid sequences of all of which we have solved by *de novo* protein sequencing with mass spectrometry. The latter technique was also used for one IgLc from a patient with AL amyloidosis. We find that all samples also contain proteases that fragment the proteins under physiologically relevant mildly acidic pH conditions, leading to amyloid fibril formation in all cases. Our results suggest that while every pathogenic IgLC has a unique ThAgg fingerprint, all sequences have comparable amyloidogenic potential. Therefore, extrinsic factors, in particular presence of, and susceptibility to, proteolytic cleavage is likely to be a strong determinant of *in vivo* aggregation behaviour. The important conclusion, which is corroborated by systematic analysis of our sequences, as well as many sequences of IgLCs from amyloidosis patients reported in the literature, challenges the current paradigm of the link between sequence and amyloid fibril formation of pathogenic light chains.

## Introduction

Protein aggregates are the hallmark, and in many cases causative agents, of severe disorders, ranging from Alzheimer’s disease to systemic amyloidoses^1–4^. The loss of protein solubility in these situations that leads to their deposition in various types of aggregates can have multiple origins, such as point mutations, post-translational modifications and over-production of proteins. The latter phenomenon occurs for example in Multiple Myeloma (MM), malignancy characterised by a clonal expansion of abnormal plasma cells in the bone marrow^5,6^. In the healthy state, they produce a complete immunoglobulin and in some cases additionally the corresponding monoclonal light chain (LC)^7^. Approximately 15% of the patients with MM produce exclusively light chains^8^. The 25 kDa sized light chain proteins are usually either excreted or degraded by the kidney, but high monoclonal quantities and low renal clearance can lead to the deposition at various sites within the kidney^9^. The deposits contain diversified kinds of IgLC aggregates, which cause different diseases, such as Light Chain Deposition Disease (LCDD), where the aggregates have an amorphous nature^10,11^, and AL-amyloidosis, where aggregates consist of amyloid fibrils^12^.

Currently, there is no possibility to predict the *in vivo* solubility and deposition behaviour of a particular monoclonal light chain found in the blood or urine of a patient. A large number of studies has been performed in recent years in order to understand the *in vivo* solubility and aggregation behaviour of a patient-derived light chain through a detailed *in vitro* investigation of the biophysical and biochemical properties^13–19^. In particular, the question as to which properties distinguish the amyloid fibril forming light chains from those that do not form amyloid fibrils in vivo has been extensively studied. Various molecular properties of the light chains, such as thermal stability^13,15,20–22^, tendency to dimerize^23–25^, or protein dynamics^26–28^ have been proposed to correlate with their deposition as amyloid fibrils. Each patient-derived light chain protein has a unique amino acid sequence determined by somatic recombination and various mutations^29,30^. The diversity of the N-terminal variable region (v-region) for recognition of the antigen^31^, is created by somatic recombination of variable (V) and joining (J) gene segments (V-J combination) during the early stages of B-cell maturation^32^. Light chains occur either as a λ or a *κ* isotype, which are encoded by the immunoglobulin λ (IGL)^33^ or the *κ* (IGK)^34^ locus. The majority of light chain amyloidosis cases are associated with λ light chains^35^. This finding clearly substantiates a correlation between sequence and the amyloid propensity of a LC. There are a multitude of different IGLV and IGKV gene segments, which are randomly joined with the corresponding joining gene segments (IGLJ and IGKJ)^36^ and then undergoing somatic hypermutations during the antigen dependent stages of differentiation. As this sequence diversity translates into a diverse clinical picture^36^ an improved understanding of the causes and mechanisms of light chain deposition requires to determine the amino acid sequences of patient-derived light chains and to correlate the sequence information with the biochemical and biophysical properties. The sequence can either be obtained through DNA sequencing of tissue from bone marrow biopsies^6,37^, or through *de novo* protein sequencing by mass spectrometry^38^. The latter, however, is challenging because of the absence of the sequence under study in the databases usually employed in mass spectrometry (MS)-based proteomics^39^. In order to investigate the link between light chain sequence and aggregation behaviour, we have developed a complete de novo protein sequencing workflow based on a combination of top-down^40^ and bottom-up proteomics with specific data analysis for patient-derived light chains. The details of this workflow are published separately^41^. The light chains investigated in this study were derived from 9 patients presenting light chains in their urine (1 isotype lambda and 8 isotype kappa) and represent a sub-set of a larger collection from a patient cohort at the University Hospital Düsseldorf, which has already been subjected to an initial biochemical and biophysical characterisation^19^. The inclusion criterion of a given sample into the present study was the availability of a sufficient amount of light chain at high purity to allow both the sequence determination, as well as the detailed Thermodynamic and Aggregation Fingerprinting (ThAgg-Fip) to be carried out. ThAgg-Fip consists of a characterisation of IgLC dimerisation, thermodynamic stability (thermal and chemical), thermally and chemically induced aggregation (amorphous and amyloid) and their respective concentration dependencies, and enzymatic digestability representing the most comprehensive characterisation of disease-related IgLCs to date. We dub our approach ThAgg-Fip, because the multi-parametric characterisation reveals a unique set of biophysical and biochemical properties of every individual patient-derived light chain. The 9 samples fulfilling our inclusion criteria are obtained from MM patients without evidence of amyloid fibril formation. Consequently our study was mostly carried out on light chain proteins, which would generally be considered to be non-amyloidogenic. However, we find that all proteins of our study are observed to form amyloid fibrils *in vitro* under physiologically relevant conditions of mildly acidic pH, where the present (co-purified) proteases led to the rapid digestion of the light chains into shorter fragments. We analyse our determined sequences as well as a representative collection of light chain sequences from the literature^42^ using several commonly used aggregation prediction algorithms and find no noticeable difference between amyloid forming and non-amyloid forming sequences. Furthermore, we were able to determine the sequence regions involved in the formation of fibrils for most of our samples and found no correspondence with predicted aggregation hot spots. Our findings challenge the current paradigm that the origin of the amyloidogenicity is to be found in the amino acid sequence of the light chains alone. We conclude that all light chains have comparable intrinsic amyloidogenicity and that the interplay between protein sequence, environmental conditions and presence and action of proteases defines whether a given light chain deposits in the form of amyloid fibrils in a given patient or not. Furthermore, our uniquely comprehensive and widely applicable ThAgg-Fip approach highlights the multiparametric nature of the problem of *in vivo* protein aggregation.

## Materials and Methods

### Patient-derived samples

The study described in this manuscript is an extension of previous work^19^ and has been reviewed and approved by the ethics committee of the University Hospital Düsseldorf. All patients of whom samples were used in the study have signed an informed consent (study number 5926R and registration ID 20170664320). Protein isolated from 24 h urine samples of 10 patients (5 females, 5 males, median age 64.5 years with a range between 45 and 72 years, 2 patients with a lambda isotype light chain and 8 patients with a kappa isotype chain) with MM and one patient with amyloidosis as detailed in table S1. A histopathological examination of the patient’s kidney was not available since the corresponding invasive diagnostic procedure was not necessary for the therapy decision-making process as they were diagnosed according to IMWG criteria. These samples represent a sub-set of samples of a previous study^19^ (the sample nomenclature is the same), and they were selected, because they contained the light chain protein at sufficiently high quantity and purity to perform the detailed biochemical and biophysical experiments and to determine the protein sequence. The protein of the patient with amyloidosis was only sequenced and not subjected to ThAgg-Fip analysis, due to lack of purity of the protein sample.

### Protein sample preparation

The IgLCs were purified as described previously^19^. Briefly summarized, the protein content of a 24 h urine collection was precipitated by ammonium sulfate (70% saturation) and the LCs were purified after dialysis by size-exclusion chromatography on an ÄKTA pure chromatography system (GE Healthcare) using a Superdex 75 10/300 GL column and 30 mM Tris-HCl, pH 7.4, as an elution buffer. IgLC concentration was determined by measuring UV-absorption at 280 nm (extinction coefficient of 33265 (P001), 27640 (P004), 26150 (P005, P006, P007, P016, P017), 33140 (P013) and 31650 M^−1^ cm^−1^ (P020). A stock solution of tris(2-carboxyethyl)phosphine (TCEP) was prepared by dissolving 100 mM TCEP in 30 mM Tris-HCl buffer pH 7.4. Subsequently, the pH value was titrated with NaOH to 7.4. Pepstatin A and E-64 were dissolved in DMSO to prepare a stock solution of 1 mM, respectively.

### LC sequence characterization and protease identification

All LC sequences were determined using a dedicated mass spectrometry workflow described in^41^. This workflow is based on the combination of bottom-up and top-down proteomics experiments with appropriate data analysis. It is important to note that the sample numbers are not identical between the present manuscript and the manuscript with the sequencing results^41^, but the latter contains a correspondence table. Briefly, light chain samples were solubilized in 8 M urea, then reduced (5 mM TCEP, 30 min) and alkylated (10 mM iodoacetamide, 30 min in the dark). After urea dilution, digestion was carried out for 3 h at 37°C (1:20) and was stopped by adding 5% FA. Peptides were desalted on Sep-Pak C18 SPE cartridge. Digests were analyzed in LC-MS/MS on a Q-Exactive Plus. Peptide elution was performed with a linear gradient on a 50 cm C18 column. Mass spectra were acquired with a Top10 data-dependent acquisition mode using classical values except for the number of MS/MS *μ* scans that was set to 4. All raw files were searched with MaxQuant against the Uniprot Homo sapiens reference proteome (74,830 entries) concatenated with the corresponding light chain sequence using trypsin as specific enzyme with a maximum of 4 miscleavages. Possible modifications included carbamidomethylation (Cys, fixed), oxidation (Met, variable) and Nter acetylation (variable). One unique peptide to the protein group was required for the protein identification. A false discovery rate cut-off of 1% was applied at the peptide and protein levels.

### Analysis of the properties of the IgLC amino acid sequences

The sequences of the IgLC samples of this study were parsed with IMGT, the international ImMuniGeneTics information system and were aligned in order to investigate the amino acid changes between the germline sequences and the sequences under study. The sequences were also analysed using different online bioinformatic tools (ZipperDB, Tango, Pasta, CamSol), which have been developed to predict the aggregation propensity/solubility of proteins based on their amino acid sequences.

### Determination of the dimeric fraction

The ratios of dimers and monomers of the various IgLCs were determined by both top-down mass spectrometry (analysis of intact proteins) and analytical ultracentrifugation (AUC). The light chains were purified by size-exclusion chromatography (SEC), aliquoted, frozen and thawed before the AUC-measurement. Prior to the MS-measurements the samples were dialyzed against ammonium bicarbonate after the SEC and freeze-dried. The chromatograms of the different samples with the fractions used for the experiments highlighted can be found in the supplementary information (supplementary figure 3). All experiments of the current study were conducted with specific fractions in order to standardise the preparation since the peaks are not always symmetrical. Therefore a bias of the measured dimer fraction due to differences in sampling cannot be fully excluded. Sedimentation velocity (SV) experiments were performed in a XL-A ProteomeLab ultracentrifuge equipped with absorbance optics. Experiments were conducted in An-60-Ti-rotor at 20°C using a rotor speed of 60,000 rpm. Solutions of 35 μM of IgLC samples were investigated and the radial scans were acquired at a wavelength which ensures an optimal resolution, which ensures an absorbance at ca. 1 OD. Scans have a radial step size of 0.002 cm, i.e. radial resolution of 500 data points/cm. The sedimentation boundaries were analysed with SEDFIT (version 16p35)^43^.

The *c*(*s*) model for sedimentation coefficient distributions as solutions to the Lamm equation implemented in SEDFIT was applied to the data. This model provides an approximation of a weight-average frictional ratio *f*/*f*_0_ for all particles of a distribution and based on which sedimentation coefficients can be translated into diffusion coefficients and/or molecular mass. An extension to the two-dimensional sedimentation coefficient distribution can be achieved with the c(*s*, *f*/*f0*) model, providing weight-average frictional ratios for each sedimentation coefficient of the distribution with color-coded signal intensities^44^.

During aggregation we observed a loss of signal due to sedimentation of large assemblies during acceleration of the centrifuge to final speed for SV analysis. Absorbance was first measured at 3,000 rpm to allow for quantification of the loss of material to 60,000 rpm. The wavelength for detection was varied to optimise the signal for SV analyses. The final *c*(*s*) curves were adjusted to have signals representative for the concentration left at the first time point of data acquisition.

The fraction of dimer measured in relation to the overall amount of native light chains (monomer and dimer) investigated by the AUC measurements was compared to the results of mass spectrometry (MS) and previously reported results from non-reducing SDS-PAGE^19^ (Figure 1).

**Figure 1.**
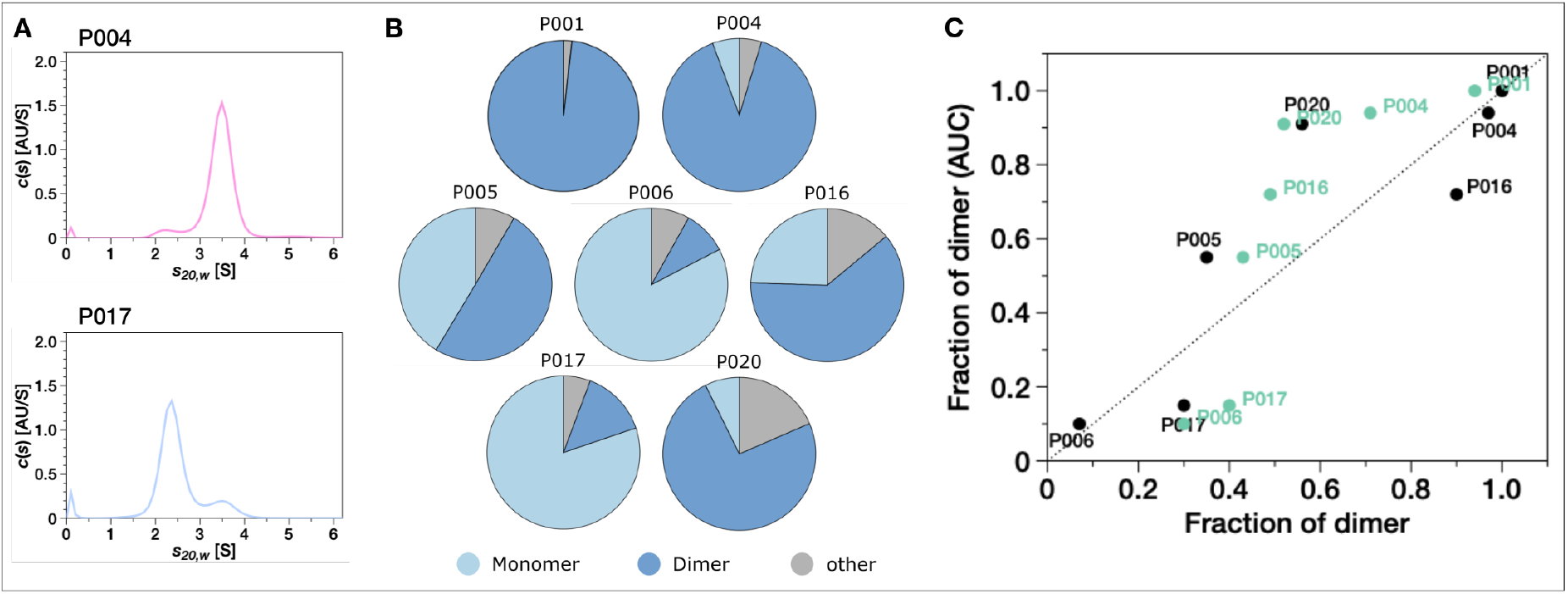
LC dimerisation (A) The distribution of sedimentation coefficients (*c*(*s*)) determined by sedimentation velocity AUC experiments at 60,000 rpm of two light chains (P004 and P017) as an example. (B) The monomer (light blue), dimer (dark blue) and other (grey) content measured by AUC. (C) The fraction of dimer measured in relation to the overall amount of native light chains (monomer and dimer) from AUC measurements against the fraction detected by MS-measurements (black) and by SDS-PAGE (green)^19^.

### Combined differential scanning fluorimetry (DSF) and dynamic light scattering (DLS) experiments

The thermal unfolding experiments of the IgLC samples as a function of protein and denaturant concentration were performed with a Prometheus Panta instrument (Nanotemper, Munich, Germany). This is a microcapillary-based (10 μl sample per capillary) instrument that allows to measure up to 48 samples in parallel. Intrinsic fluorescence can be excited at 280 nm and emission is monitored at 330 nm and 350 nm. The DLS experiments are performed on the same sample with a laser at 405 nm. Furthermore, the instrument also allows to measure the sample turbidity with back reflection optics. The experiments where the initial increase and subsequent decrease in turbidity of the samples was followed (‘Bence-Jones test’, BJT) were performed by scanning the temperature from 25 to 90°C at a scan rate of 2.5°C/min. The other thermal ramping experiments were performed by scanning from 20°C (urea dependence) or 25°C (concentration dependence) to 70°C at a scan rate of 1°C/min. For the melting scans, we prepared stock solutions of the IgLCs (97 μM (P001), 150 μM (P004), 91 μM (P005), 139 μM (P006), 103 μM (P007), 97 μM (P013), 99 μM (P016), 158 μM (P017) and 81 μM (P020)). These stock solutions were filtered through a 220 nm pore size syringe filter before concentration determination and the DSF-DLS experiments. The BJT was performed at a uniform sample concentration of 30μM and at pH 5.0, which was established by adding 100 mM Na-acetate buffer to the stock solutions in 30 mM Tris buffer pH 7.4. A pH value of 5 was used in the BJT as this corresponds to the original protocol used to test patient urine for BJ-proteins^45^. For the concentration-dependent measurements, the stock solutions were diluted 3 times by a factor of two with 30 mM Tris buffer pH 7.4, to yield 4 different concentrations per protein, allowing to define the concentration-dependence of the unfolding and aggregation temperatures. For the urea-dependent experiments, the stock solutions were diluted 5-fold into solutions of appropriate urea concentration. The final urea concentrations were 0, 0.67, 1.34, 2.01, 2.68, 3.35, 4.02, 4.69, 5.36 M and the buffer concentration was in all cases 30 mM Tris buffer. The data was visualised as the ratio of the intrinsic fluorescence emission intensity at 350 nm over the intensity at 330 nm. For the thermal unfolding at the different protein concentrations, the melting temperature (T_*m*_) and the temperature of aggregation onset (T_*agg*_), as well as the cumulant radius, were automatically determined by the instrument software. The melting temperature corresponds to the maximum of the first derivative of the change in fluorescence intensity emission ratio. The onset of aggregation is defined by fitting the cumulant radius as a function of temperature by both a linear function (i.e. an extrapolation of the baseline) and by a sigmoidal model. The onset of aggregation is defined as the temperature at which the two fits first differ by more than 0.5 %. For the experiments in the absence of denaturant, the buffer viscosity and its temperature dependence was set to that of water. In the samples with urea, we did not correct the viscosity, which was not necessary as we did not analyse the sizes precisely. In these experiments, DLS was merely used to determine which samples had formed aggregates, and should therefore be excluded from the thermodynamic analysis. In Table 1 we report the values for T_*m*_ (pH 5 and 7.4) and T_*agg*_ (pH 7.4), as well as the concentration dependencies, 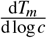 and 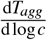 (pH 7.4). We use the logarithmic derivatives, because the dependency is approximately linear on a logarithmic concentration scale, which allows to use the slope of a linear fit as a single key parameter across the entire concentration range explored. It has been shown that the determination of the unfolding temperature from fluorescence intensity ratios can lead to systematic deviations, depending on the relative intensities of the fluorescence spectra of the folded and unfolded states^46^. The use of the derivative introduced above eliminates such systematic errors, because a constant offset of all melting temperatures does not affect their dependence on protein concentration. We also visualise the evolution of the size distribution of the sample as a function of temperature with a heat map on a logarithmic size scale. The full sets of raw data of these experiments can be found in supplementary figures 5 and 6. In these plots, each time/temperature point corresponds to a full particle size distribution determined from a multi-species fit to the intensity autocorrelation function. These plots are automatically generated by the control software of the Prometheus Panta instrument. For the combined chemical and thermal unfolding experiments, the data set of each protein was globally fitted to the thermodynamic two-state model recently presented^47^. The global fits are shown in supplementary figure 7. In order to reduce the influence of aggregation on the fits, only samples containing urea were included in the fit, as the simultaneous DLS measurements had shown aggregation mostly in the absence of urea. For P004, all samples below 2 M urea were excluded on this rationale. From the global fit over all temperatures, we then determine the stability of the IgLC, ΔG, at 37°C. We find a significant correlation between ΔG and the m-value, i.e. 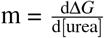^48^. We therefore fix the m-value to a common value for all different light chains and focus on the resulting differences between ΔG. Error estimates of the obtained values were obtained by 100-fold bootstrapping by resampling the different capillaries with replacement.

**Table 1.** Overview over the thermodynamic and aggregation fingerprints of the IgLCs of this study. All T_m_ values, turbidity maxima and aggregate sizes are reported at a concentration of ca. 30 μM. The DSF experiments at pH 5 (pH 7.4) were performed at a scan rate of 2.5°C/min (1.0°C/min). The concentration dependence of the melting and aggregation temperatures was explored in the range between ca. 15 and 120 μM. *m-value fixed for all samples. **Different m-value allowed in fit. ***Only one population with intermediate *s*-value detected. ✓ThT-positive aggregation observed.

|  | P001 | P004 | P005 | P006 | P007 | P013 | P016 | P017 | P020 |
| --- | --- | --- | --- | --- | --- | --- | --- | --- | --- |
| dimerisation by SDS-PAGE | 0.96 | 0.71 | 0.43 | 0.3 | 0.32 | 0.36 | 0.49 | 0.4 | 0.52 |
| dimerisation by MS | 1 | 0.97 | 0.35 | 0.07 |  |  | 0.9 | 0.3 | 0.56 |
| dimerisation by AUC | 1 | 0.94 | 0.55 | 0.1 | *** | *** | 0.72 | 0.15 | 0.91 |
| $T_m$ DSF (pH 5) [°C] | 63.7 | 51.1 | 51.0 | 54.9 | 50.3 | 51.7 | 54.1 | 50.4 | 56.3 |
| T (max. turb. @ pH 5) [°C] | > 90 | 62 | 71 | no turb. | 66 | 64 | no turb. | 65 | 83 |
| $T_m$ DSC (pH 7.4) [°C] <sup>19</sup> | 64.6 | 58.5 | 51.2 | 50.4 | 55.4 | 52.2 | 56.8 | 51.7 | 53.5 |
| $T_m$ DSF (pH 7.4) [°C] | 62.6 | 54.2 | 50.9 | 50.6 | 53.8 | 52.6 | 54.7 | 49.4 | 50.4 |
| d $T_m$ / d log(c) (pH 7.4) | -0.33 | -1.19 | -1.20 | 0.9 | -0.55 | -1.10 | -0.17 | 0.15 | 0.23 |
| $T_{agg}$ DSF (pH 7.4) [°C] | 54.9 | 53.0 | 50.8 | 59.4 | 55.2 | 50.2 | n/a | 61.7 | 50.3 |
| d $T_{agg}$ / d log(c) (pH 7.4) | -5.8 | -2.9 | -4.1 | -1.5 | -3.2 | -4.1 | -0.5 | -22.0 | -4.6 |
| Cumul. rad. (pH 7.4, 70°C) [nm] | >1000 | >1000 | ~1000 | 7 | ~300 | ~25 | 5.5 | 16 | >1000 |
| Fract. refold. - DSF | small | small | small | large | small | small | medium | large | small |
| Fract. refold. - DSC <sup>19</sup> | 0 | 0.18 | 0.36 | 0.68 | 0.33 | 0.38 | 0.41 | 0.7 | 0.22 |
| $\Delta G$ [kJ/mol] at 37°C | 26.3±1.5 | 25.3±0.4 | 20.8±2.1 | 20.8±0.8 | 23.7±0.6 | 21.2±0.7 | 10.0±0.7 | 21.8±1.1 | 20.2±0.5 |
| m-value | 7.1* | 7.1* | 7.1* | 7.1* | 7.1* | 7.1* | 2.3±0.5** | 7.1* | 7.1* |
| digestability by trypsin <sup>19</sup> | 0.1 | 1 | 0.7 | 0.24 | 1 | 1 | 0.17 | 0.98 | 1 |
| acid induced amyloid form. | ✓ | ✓ | ✓ | ✓ | ✓ | ✓ | ✓ | ✓ | ✓ |

### Measurements of aggregation kinetics

Different solution conditions (acidic pH values) were tested for their potential to induce aggregation of patient-derived, purified IgLCs. In order to prepare the samples at different acidic pH values (pH 2, pH 3, pH 4), protein solutions of different concentrations were diluted from 30 mM Tris-HCl pH 7.4 1:1 into 300 mM citric acid buffer at the desired pH value). Two or three replicates of each solution were then pipetted into a high-binding surface plate (Corning #3601, Corning, NY, USA). The aggregation kinetics were monitored in the presence and absence of small glass beads (SiLibeads Typ M, 3.0 mm). The plates were sealed using SealPlate film (Sigma-Aldrich #Z369667). The kinetics of amyloid fibril formation were monitored at 37°C either under continuous shaking (600 rpm) or under quiescent conditions by measuring ThT fluorescence intensity through the bottom of the plate using a FLUOstar (BMG LABTECH, Germany) microplate reader (readings were taken every 150 or 300 seconds). In order to compare the factor of the increase of the ThT fluorescence emission intensity between the samples, the ThT fluorescence emission intensity at the end of the experiment was compared with the lowest emission value. The halftimes of the aggregation reaction are defined as the point where the ThT intensity is halfway between the initial baseline and the final plateau. The halftimes were obtained by individually fitting the curves using the following generic sigmoidal equation^49^:

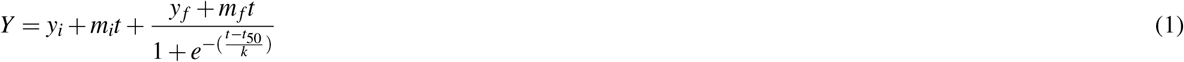

Where *Y* is the ThT fluorescence emission intensity, *t* is the time and *t*_50_ is the time when 50 % of maximum ThT fluorescence intensity is reached. The initial baseline is described by *y_i_* +*m_i_t* and the final baseline is described by *y_i_* +*m_i_t*+*y_f_*+*m_f_t*. While this equation does not describe the underlying molecular processes of aggregation, it does allow determination of the main phenomenological parameter, the half time, of each experiment. At the end of the experiments, the amount of aggregated protein was determined by combining the replicates and pelleting the aggregation product for 1 h at 16,100 g followed by measuring the soluble content by UV-absorbance at 280 nm of the supernatant, and correcting for the absorbance of the ThT. In order to investigate whether the aggregation can be seeded at acidic pH, fibrils were produced by incubating 35 μM protein solution at pH 3 and pH 4 in a high-binding plate and in a 2 ml Eppendorf tube in presence of glass beads under shaking conditions at 37°C. The presence of fibrillar aggregates was confirmed by AFM. For seed generation the fibril solutions obtained from the plate and from an Eppendorf tube were homogenized using an ultra-sonication bath Sonorex RK 100 H (Bandelin, Germany) for 300 s.The seeded aggregation experiments were performed in high-binding surface plates under quiescent conditions with 35 μM P016 monomer and the preformed seeds were added to a final concentration of 5 % of the monomer solution at the desired pH value. The seeds were added either at the beginning or after 6 h pre-incubation of the monomer solution at 37°C. We furthermore investigated the seeding potential at neutral pH, where the protein remains largely unaffected by proteolytic activity. We used pre-formed fibrils, which were produced at pH 3 and pH 4 in an Eppendorf tube as described above at 100 μM LC concentration. The seeds were additionally washed to remove the soluble fragments by centrifuging the sample at 137,000 g at 20°C for 45 min and resuspending the pellet in 150 mM citric acid. This washing procedure was carried out three times. The seeded aggregation experiments were performed in high-binding surface plates under agitation conditions with 50 μM monomer solution and seeds added to a final concentration of 5 %.

### Prevention of amyloid fibril formation at acidic pH values

In order to investigate whether the cleavage of the IgLCs at acidic pH values was responsible for the amyloid fibril formation, we tested whether inhibiting some of the identified proteases will prevent the formation of ThT-positive aggregates. Therefore the IgLCs (35 μM monomer concentration) were incubated as described above in a high binding surface plate under quiescent conditions in the presence of 10 μM pepstatin A and E-64, respectively. E-64 is an irreversible and highly selective cysteine protease inhibitor, pepstatin A is a reversible inhibitor of acidic proteases (aspartic proteases) and can be used in a mixture with other enzyme inhibitors. Further experiments were conducted with P005 and P016 in the presence of 1 μM pepstatin A or E-64 or 1 μM or 10 μM pepstatin A and E-64 under agitation conditions. The morphology of aggregates was investigated by AFM.

### Fibril fragment determination

The aggregation products formed in Eppendorf tubes under agitation conditions at pH 3 and pH 4 were centrifuged using an Optima MAX-XP ultracentrifuge (Beckman Coulter) in a TLA-55 rotor at 40,000 rpm at 20°C for 45 min. The pellet was re-suspended in 150 mM citric acid (pH 3 or pH 4) and centrifuged again for 45 min. This washing procedure to remove the soluble fragments was conducted three times. The washed aggregates were dissolved in 6 M urea and subsequently analysed by mass spectrometry.

A Dionex UltiMate 3000 RSLC Nano System coupled to an Orbitrap Fusion Lumos mass spectrometer fitted with a nano-electrospray ionization source (Thermo-Scientific) was used for all experiments. 5 μL of reduced/alkylated protein samples in solvent A were loaded at a flow rate of 5 μL min-1 onto an in-house packed C4 (5 μm, Reprosil) trap column (0.150 mm i.d. x 35 mm) and separated at a flow rate of 0.5 μL min-1 using a C4 (5 μm, Reprosil) column (0.075 mm i.d. x 28 cm). The following gradient was used: 2.0% B from 0–10 min; 20% B at 11 min.; 60% B at 22 min.; 99% B from 25-30 min.; and 2.0% B from 30.1 to 35 min. Solvent A consisted of (98% H2O, 2% ACN, 0.1% FA) and solvent B (20% H2O, 80% ACN, 0.1% FA). MS scans were acquired at 120,000 resolving power (at m/z 400) with a scan range set to 550–2,000 m/z, four microscans (μscans) per MS scan, an automatic gain control (AGC) target value of 5×105 and maximum injection time of 50 ms. MS/MS scans were acquired using the Data-Dependent Acquisition mode (Top 4) at 120,000 resolving power (at m/z 400) with an isolation width of 1.2 m/z, five μscans, an AGC target value of 5×105 and maximum injection time of 250 ms. For fragmentation, electron transfer dissociation with 10 ms of reaction injection time and a supplemental higher-energy collisional dissociation with normalised collision energy (NCE) of 10% (EThcD) was used. All data were processed with ProSightPC v4.1 (Thermo-Scientific) and Proteome Discoverer v2.4 (Thermo-Scientific) using the ProSightPD 3.0 node. Spectral data were first deconvoluted and deisotoped using the cRAWler algorithm. Spectra were then searched using a two-tier search tree with searches against the corresponding LC sequences. The search 1 consists of a ProSight Absolute Mass search with MS1 tolerance of 10 ppm and MS2 tolerance of 5 ppm. The search 2 is a ProSight Biomarker search with MS1 tolerance of 10 ppm and MS2 tolerance of 5 ppm. Identifications with E-values better than 1e-10 (-log (E-value) = 10) were considered as confident hits.

### Atomic Force Microscopy (AFM)

Atomic force microscopy height-images were acquired after the aggregation kinetic measurements. 10 μL of each sample (after diluting 1:4 with dH_2_O) was deposited onto freshly cleaved mica. After drying, the samples were washed 5 times with 100 μL of dH_2_O and dried under gentle flow of nitrogen. AFM images were obtained using a NanoScope V (Bruker) atomic force microscope equipped with a silicon cantilever ScanAsyst-Air with a tip radius of 2-12 nm. The images were analyzed with the software Gwyddion 2.56 to measure height profiles and investigate a possible twisting of the fibrillar aggregates.

### Microfluidic diffusional sizing and concentration measurements

Fluidity One is a microfluidic diffusional sizing (MDS^50^) device, which measures the rate of diffusion of protein species under steady state laminar flow and determines the average particle size from the overall diffusion coefficient. The protein concentration is determined by fluorescence intensity, as the protein is mixed with ortho-phthalaldehyde (OPA) after the diffusion, a compound which reacts with primary amines, producing a fluorescent compound^51^. To measure the influence of pH on the average size of the molecules in the solution, the protein was pre-incubated at acidic pH values with 150 mM citric acid (pH 3 or pH 4). The IgLC solution was incubated in an Eppendorf tube at 37°C under quiescent conditions. After different incubation times, 6 μL of the solutions were pipetted onto a disposable microfluidic chip and measured with the Fluidity One (F1, Fluidic Analytics, Cambridge, UK).

### Statistical analysis

To investigate the correlations between the experimental quantities measured of the IgLCs, numerical values were needed for all measurements. This necessitated an arbitrary conversion of fraction refold by DSF to a numerical scale of 0.33, 0.50, or 0.66, for small, medium, and large, respectively. The ΔG-value was not included for P016, and m-values were not used because they were globally fitted and therefore shared between all data sets. All other non-numerical cells in Table 1 were interpreted as missing values. Before training an Elastic Net model on the data, missing values were set to the mean of the given observable and all values were normalised to the mean value with unit variance. The following limited set of parameters were chosen to eliminate the most internally correlated measurements (e.g. dimerization by three different methods): *T_m_* DSF (pH 5), T (maximum turbidity at pH 5), *T_m_* DSC, 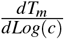 (pH 7.4), 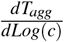 (pH 7.4), Cumulant radius (pH 7.4, 70°C), Dimerization by AUC, Fraction refold by DSC, dG (37°C), and Digestability by trypsin. Model training was done using a grid search with 4-fold cross validation using negative mean squared error as the loss function. A separate validation set was not created because of the low number of data points. This analysis was performed in Python 3.7 using packages Numpy, Pandas, SKLearn, Matplotlib and Seaborn.

## Results

We set out to perform a comprehensive characterisation of the thermodynamic and aggregation properties of the patient-derived light chains with the aim to combine this information with the sequences that we obtained from a parallel sequencing effort^41^. To this end, we have combined a wide array of state of the art biophysical and biochemical methods into the ThAgg-Fip approach. We included the experimental parameters that have previously been reported to correlate with light chain amyloidogenicity, such as dimerisation, thermal stability and proteolytic digestability. Furthermore, we included a detailed characterisation of the thermodynamics and aggregation behaviour of each light chain. The resulting ThAgg fingerprints are listed in table 1.

### Dimerisation of the light chains

We first set out to robustly characterise the monomer-dimer distribution of the IgLCs. The tendency of disease-related overproduced immunoglobulin light chains to dimerise has previously been proposed to correlate with the protein’s tendency to form amyloid fibrils^23,25^. In a previous study, we determined the degree of dimerisation with non-reducing SDS-PAGE gels^19^. Here we aimed to improve our previous results by both sedimentation velocity analytical ultracentrifugation (AUC), as well as mass spectrometry. AUC allows the analysis under native conditions in solution and mass spectrometry allows unique identification of protein monomers and multimers. A detailed discussion of the dimerisation behaviour, including the observation of mixed dimers formed from two IgLC isoforms, can be found in^41^. The fractions of monomer and dimer displayed in Figure 1 A are extracted from sedimentation velocity *c*(*s*) profiles. Both species could be fitted as two distinct distributions in the displayed samples. As ‘other’, we denote additional contents of some of the samples, for example HSA (in P020) or aggregates. The measured average *s*_2_0, *w*-value for the monomer is 2.32±0.07 S and for the dimer 3.53±0.18 S. The *c*(*s*) profiles of P007 and P013 displayed only one species situated between the s-value of a monomer and a dimer. We also find that the dimers can differ surprisingly strongly in their form factors, ranging from close to globular to elongated (see supplementary figure 4). Taken together, these techniques provide a very accurate picture of the monomer-dimer distribution of the light chain proteins in solution. None of the examined IgLCs occurs exclusively in its monomeric form, but P001 is exclusively dimeric. Overall, our samples span a wide range of degree of dimerisation (~0.1-1). The results of the three different methods are displayed in Figure 1 C and listed in table 1. The three techniques exhibit similar outcomes. Especially P001, P004 and P006 behaved very similar in the AUC- and the MS-measurement, despite the different sample treatments before the measurements (see methods).

### Thermal and chemical stability and thermally induced aggregation

One of the most intriguing and defining features of IgLCs is their temperature-induced aggregation behaviour at mildly acidic pH (~5). Upon heating to ca. 60°C, IgLC solutions become turbid, and upon further heating to above 90°C, the turbidity is partly reversible. This property, which is not well understood, was historically used to detect free IgLCs in urine^45^. This method was eventually replaced by more modern methods, because of the limited reliability. We simulated this method by performing rapid thermal ramping experiments from 25 to 90°C (see methods section for details) and found indeed that the majority of the samples showed a decrease of turbidity above ca. 70°C (Figure 2 a). However, the turbidity, its temperature maximum and degree of its reversibility varied greatly between the samples. We then proceeded to characterise the thermal and chemical stability as well as aggregation behaviour of the IgLC samples at neutral pH (7.4). We performed thermal ramping experiments (25-70°C) at different protein concentrations (see methods) while simultaneously measuring the intrinsic Trp fluorescence of the IgLC samples as well as performing dynamic light scattering (DLS) experiments. An example of such experiments (for P007) is shown in Figure 2 b. At the beginning of the experiment, the native protein (monomeric and dimeric) dominates the DLS signal in all cases. We found a dependence of both the unfolding temperature, as well as the temperature of onset of aggregation, on the protein concentration, and we quantify these dependencies by defining the parameters 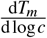 and 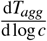.

**Figure 2.**
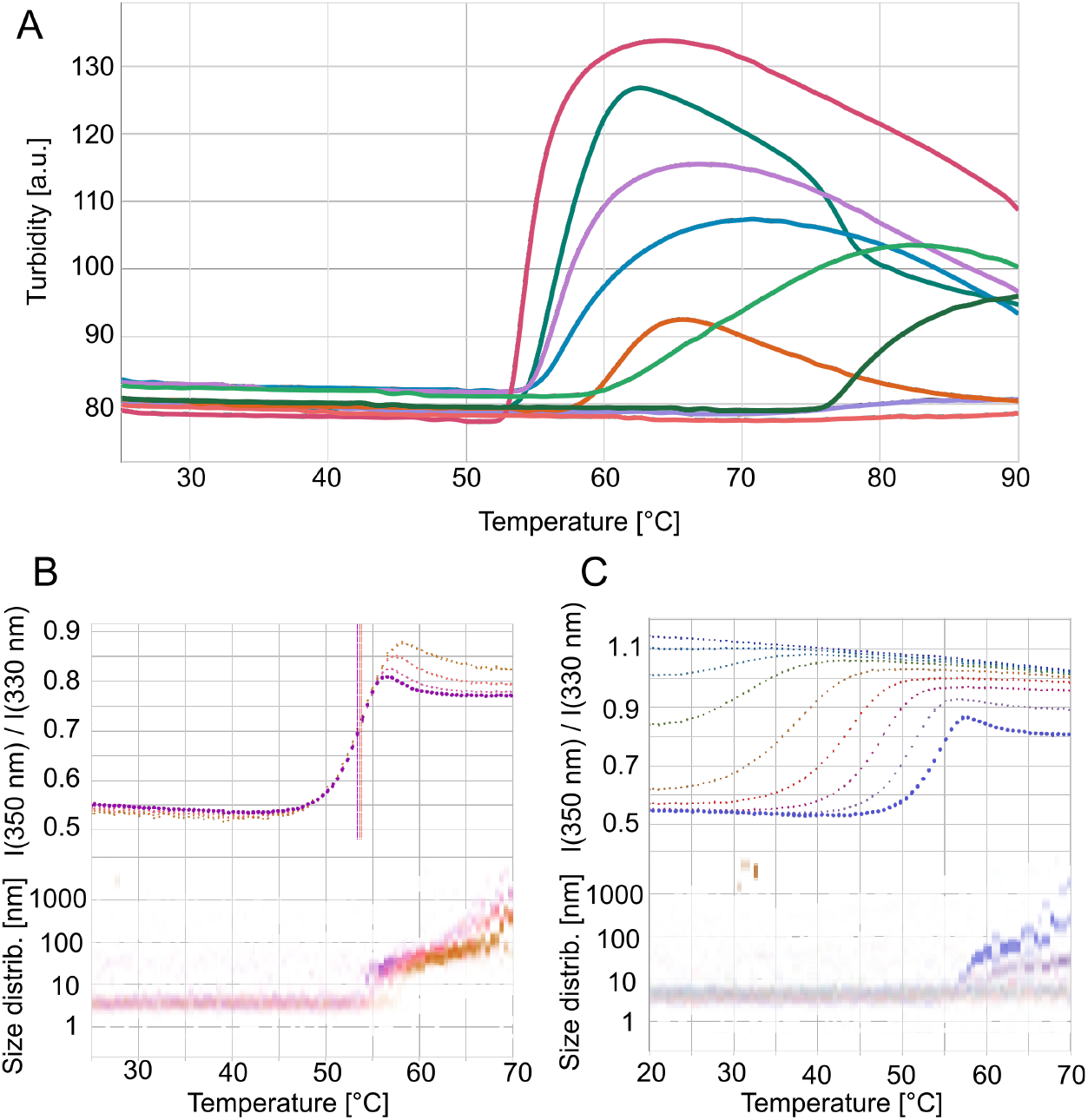
Thermal stability and aggregation of IgLCs. (A) Sample turbidity is measured as IgLC samples at pH 5 are heated from 25 to 90°C. (B) Thermal unfolding of P007 (as an example) at 103, 52, 26 and 13 μM followed by intrinsic fluorescence. In addition, the evolution of the size distribution is shown as a function of temperature (full data set with all samples in supplementary figure 5). (C) Combined chemical and thermal unfolding of P007 (as an example) at 20 μM and urea concentrations of 0 to 5.36 M in 8 steps (full data set with all samples in supplementary figure 6). The fit to the raw data can be found in supplementary figure 7, and the temperature dependent chemical stability curves in supplementary figure 8.

While 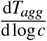 is always negative, we found that 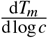 can be both negative and positive. It has been reported that aggregation induced by unfolding can lead to a concentration-dependent decrease in unfolding temperature according to the law of mass action^52^, whereas molecular crowding can have the opposite effect, i.e. a concentration-dependent stabilisation^53^. The onset and degree of aggregation vary considerably between the samples. Aggregation can start significantly before (e.g. P001) or after (e.g. P006) the midpoint of unfolding. Furthermore, the aggregate sizes (as quantified from the cumulant radius of the samples at 70°C) vary between sizes below 10 nm (e.g. P016) and sizes of more than 1 μm (e.g. P020). We also followed the degree of re-folding by cooling the samples down and found that most samples (except P006 and P017) show very little re-folding.

Next, we extended these measurements by performing a temperature ramp of samples at different denaturant (urea) concentrations; examples are shown in Figure 2 c, where the fluorescence ratio (350 nm/330 nm) and the DLS size distributions are shown. As expected, the proteins become increasingly destabilised at higher temperatures. Interestingly, we find that significant aggregation is only observed in the absence of urea, as well as for the 1-2 lowest urea concentrations. This information allowed us to perform a global fit to a model of the temperature-dependent IgLC thermodynamic stability^47^, where we excluded the samples where aggregation was observed. Details of the modelling, which was performed directly on the fluorescence intensity rather than the ratio, can be found in the methods section. All the global fits can be found in supplementary figure 7. While this analysis allows to define the stability at any temperature in the measured interval, we report only the folding free energy, ΔG, at 37°C as a measure for the protein stability under physiological conditions in table 1, together with the other data on thermal stability and aggregation discussed above.

### LC aggregation at acidic pH values

In our previous study we found that some of the investigated light chains formed amyloid fibrils at mildly acidic pH values (pH 4), even though they came from patients without confirmed AL amyloidosis^19^. As we had detected a strong influence of the nature of the reaction vessel surface on the aggregation kinetics, we decided to examine the aggregation behaviour of the IgLC in the present study more systematically, at pH 2, pH 3 and pH 4 in high-binding polystyrene multi-well plates (Figure 3). All samples and conditions displayed an increase in ThT fluorescence intensity, except P001 and P017 at pH 4, but the relative increase in the fluorescence intensity differs significantly between the samples. The most efficient aggregation kinetics was detected at pH 3. In Figure 3 G, AFM height images of products from the aggregation experiment at pH 2 are shown. The detected amyloid fibrils appeared very distinct in the different samples: as mature short fibrils (P004), as mature long fibrils, which are associated with amorphous aggregates (P006), as a mixture of mature and single-stranded fibrils (P005) or as mainly single-stranded and protofibrils (P013, P020). Additional AFM images of fibrils formed in different reaction vessels can be found in supplementary figure 10.

**Figure 3.**
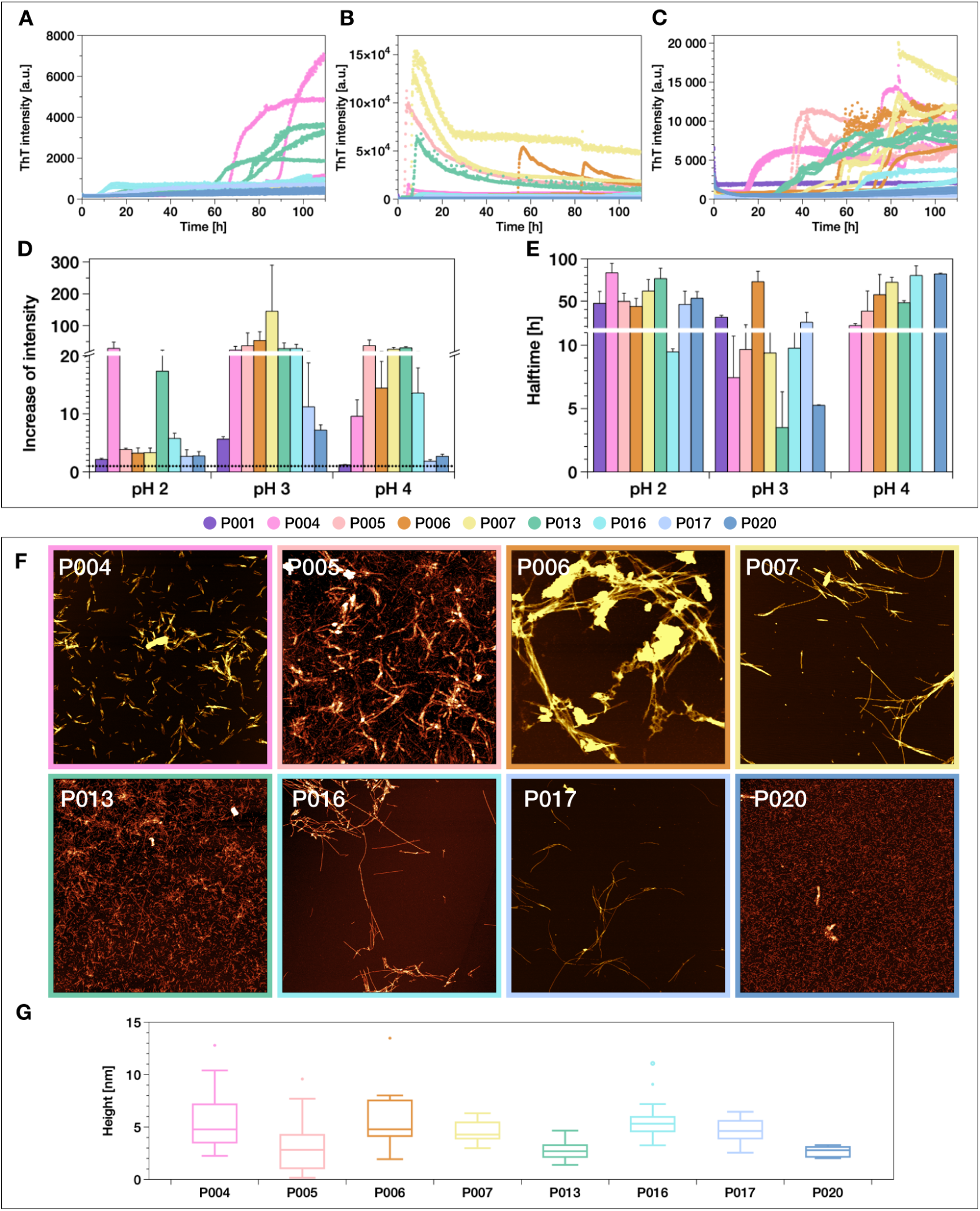
ThT fluorescence aggregation assay of the IgLC samples at (A) pH 2, (B) pH 3 and (C) pH 4 monitored in high-binding plates in the presence of glass beads under conditions of mechanical agitation (top). The aggregation kinetics are analysed by the (D) increase of intensity and (E) aggregation halftime. (F) AFM-height-images of IgLC samples after aggregation at pH 2. The image scale is 5 × 5 μm. The colour range represents the height from −2 to 15 nm, (G) box plot of the observed values of the height is illustrated.

Since the aggregation at pH 3 is the most rapid we examined the aggregation behaviour of the IgLCs at different monomer concentrations (5 μM, 35 μM and 100 μM) under this pH condition (Figure 4). The plate was shaken in the absence of glass beads. Apart from P001, whose aggregation had no lag time, the aggregation kinetics followed the typical sigmoidal time course of amyloid formation at first glance. The lag phase can last for several hours, but as soon as an increase in ThT fluorescence was detected, the aggregation reaction was very fast. After reaching the steady-state phase, the ThT-signal was found to decrease in the case of the samples at 100 μM concentrations. This decrease can be explained by the formation of insoluble, big aggregates, which sediment and disappear out of the focus of detection. The relative increase in fluorescence intensity was found to be very substantial in some cases; for example for P004 and P007 the intensity increased by a factor of over 1000. P004 aggregated as one of the fastest at the concentrations of 35 μM and 100 μM, however showed a twofold longer halftime at 5 μM. The aggregation process of some samples (P005, P013, P016 and P020) was found to be more complex, because it features a bi- or triphasic behaviour. We tested the influence of glass beads and preformed fibrils (seeds) on the aggregation kinetics of P016 in more detail as an example (supplementary figure 9). We pre-incubated the protein (35 μM monomer concentration) at pH 3 and pH 4 in a high-binding surface plate and added glass beads after 24 h or seeds after 6 h. The glass beads which were added after the pre-incubation had less of an accelerating effect on the amyloid formation compared to glass beads added at the start of the aggregation assay (pH 4) or merely shaking (pH 3). We found that the amyloid fibril formation could be seeded by adding seeds formed under different conditions. However the amyloid kinetics were not typical for a pronounced seeding effect, especially at pH 4. At this condition, even though the seeds accelerated the fibril formation significantly, a lag time was still observed. If the proteins were pre-incubated at this mildly acidic pH, on the other hand, rapid aggregation was observed from the moment of addition of the seeds at pH 3.

**Figure 4.**
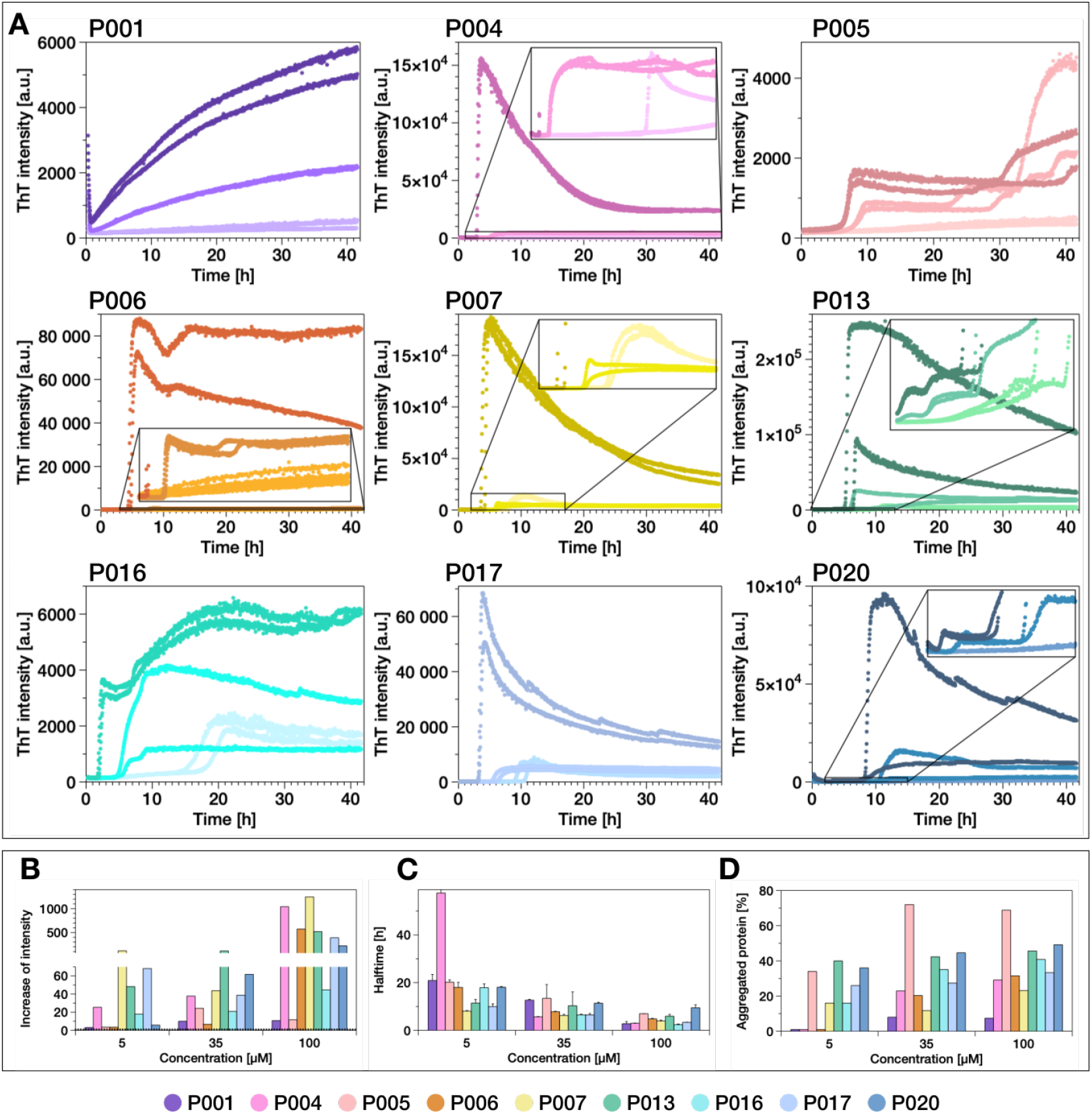
(A) ThT fluorescence aggregation assay of the IgLC samples at pH 3 in the presence of glass beads measured in high-binding surface plate under agitation conditions. The different colour shades indicate the different monomer concentrations: light: 5 μM; intermediate: 35 μM; darK: 100 μM. Overview of the effect of pH 3 on IgLC aggregation assayed by (B) increase of ThT fluorescence intensity, (C) the halftime of the aggregation and (D) relative fraction of aggregated protein determined by measuring the remaining soluble content after centrifuging the end product of the aggregation reactions. The three replicates per condition were combined before centrifugation.

The influence of the pre-incubation of the monomer at acidic pH suggests a possible modification that the monomeric proteins might undergo under these conditions. We therefore set out to probe whether the proteins can undergo proteolysis. For this purpose the samples were incubated in an Eppendorf tube at 37°C under quiescent conditions, in order to slow down as much as possible the formation of amyloid fibrils. The samples were analysed at different time points using SDS-PAGE and the microfluidic diffusional sizing^50^ device Fluidity One (Figure 5). The time course of the relative proportions of native IgLCs (monomer and dimer combined) was determined by SDS-PAGE. Incubation at pH 3 was found to have a strong effect on the size of the monomeric protein; already after one hour incubation, almost no full length protein was found to remain. Only P001, which occurs exclusively in dimeric form, partly resisted acid induced degradation over several hours. At pH 4 the samples were found to be more resistant, but after an incubation of 24 h the IgLCs were found to be fragmented to between 50 and 100 percent. The fragmentation at pH 2 was found to be faster than that at pH 4, but slower than at pH 3. P013 and particular P001 were found to be comparatively more resistant against the cleavage at pH 2. The degradation of the IgLCs at pH 3 and pH 4 could also be observed with the Fluidity One instrument, which allows to measure the average radius of the sample molecules and the protein concentration. At pH 3 the radius decreased in the first hour by about 1 nm and the IgLC samples have an average radius of around 1 nm after 160 h, which equals the average radius at pH 4 after this incubation duration. P013 appeared stable at pH 4 regarding the hydrodynamic radius, but the measured concentration displayed a strong decrease, which can possibly be explained through the formation of aggregates. Amorphous aggregates as well as fibrils are not detected in the same quantitative manner by the Fluidity One instrument, because large aggregates may not be able to enter the microfluidic channels. Furthermore, aggregates may not be as efficiently stained by the fluorescent modification used for protein quantification in the Fluidity One (Figure 5B).

**Figure 5.**
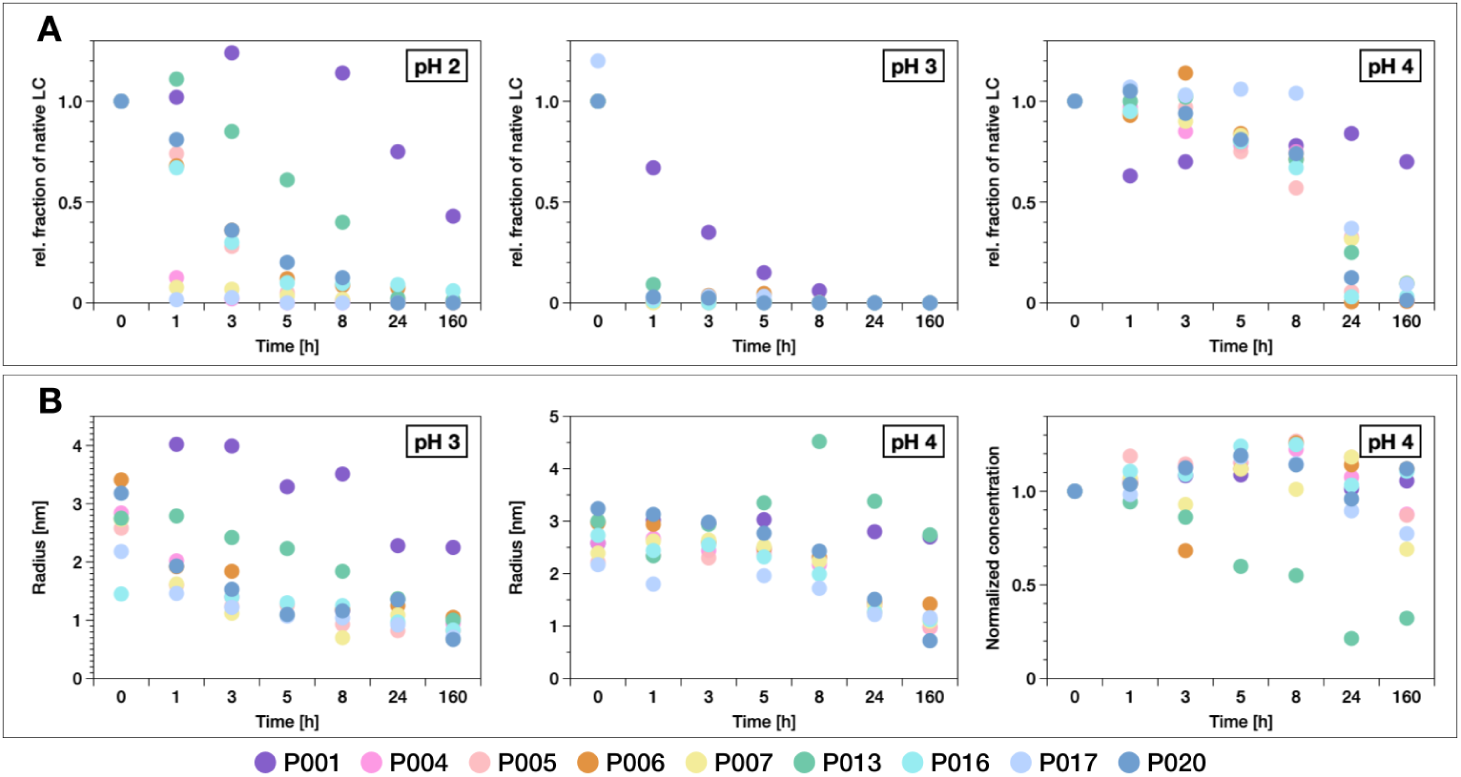
The influence of acidic pH on the IgLC samples. (A) The fraction of native protein (monomer and dimer combined) at different incubation times determined by SDS-PAGE (left: pH 2, middle: pH 3, right: pH 4) and (B) the hydrodynamic radius in nm (left: pH 3, middle: pH 4) and the normalized concentration measured by FluidityOne (right: pH 4).

Such an efficient fragmentation of the IgLCs at mildly acidic pH and room temperature cannot easily be explained by acid-catalysed hydrolysis of the polypeptide backbone. We therefore searched for the possible presence of proteases using a regular MS-based proteomics approach. The results indeed revealed the presence of different cathepsins in the IgLC samples purified from patients urine (see Table S2). Cathepsins are predominantly endopeptidases, which are located intracellularly in endolysosomal vesicles, but can furthermore be found in the extracellular space. They are important regulators and signalling molecules of various biological processes and are involved in the production of inflammatory cytokines and enhancement of tumour development^54,55^. The activity of cysteine cathepsins is increased at slightly acidic pH values and they are mostly unstable at neutral pH^55,56^. In order to probe whether the fragmentation of the IgLC at acidic pH values was caused by the present cathepsins, we incubated the samples with protease inhibitors. As inhibitors we choose E-64 and pepstatin A. E-64 (trans-Epoxysuccinyl-L-leucylamido(4-guanidino)butane) is an active-site directed, irreversible inhibitor of cysteine proteases and is known to inhibit cathepsin B, cathepsin L, cathepsin H and cathepsin Z^57–61^. Pepstatin A (isocaleryl-L-valyl-L-valyl-4-amino-3-hydroxy-6-methylheptanoyl-L-alanyl-4-amino-3-hydroxy-6-methylheptanoic-acid) is a very selective and potent inhibitor of cathepsin D, which is one of the major aspartyl endopeptidases in mammalian cells and has an pH-optimum at 3.5-5^62^. Cathepsin D was the second most frequent cathepsin present in the samples (in P017 it had the highest fraction)^63^.

The IgLCs (35 μM monomer concentration) were incubated at pH 3 in the presence of 10 μM pepstatin A and 10 μM E-64 in a high-binding plate and the potential aggregation was followed by the increase of ThT-fluorescence intensity (Figure 6). With the exception of P005 and P006, all samples showed an increase in ThT-signal. However, the ThT intensity was found to increase only by a factor of approximately two, which is almost negligible compared to the increase observed in the absence of protease inhibitors. AFM-imaging of the samples displayed small, globular oligomeric structures, very different from the mature aggregates observed to form in the absence of the protease-inhibitors. P006 and P016, which did not or slowly aggregate, showed slightly elongated aggregates, which may be pre-fibrillar structures. While the incubation of the samples at pH 3 led to a complete fragmentation of the native IgLCs, the presence of the inhibitors was found to maintain the IgLCs in their original size, even after incubation for 50 h (Figure 6C).

**Figure 6.**
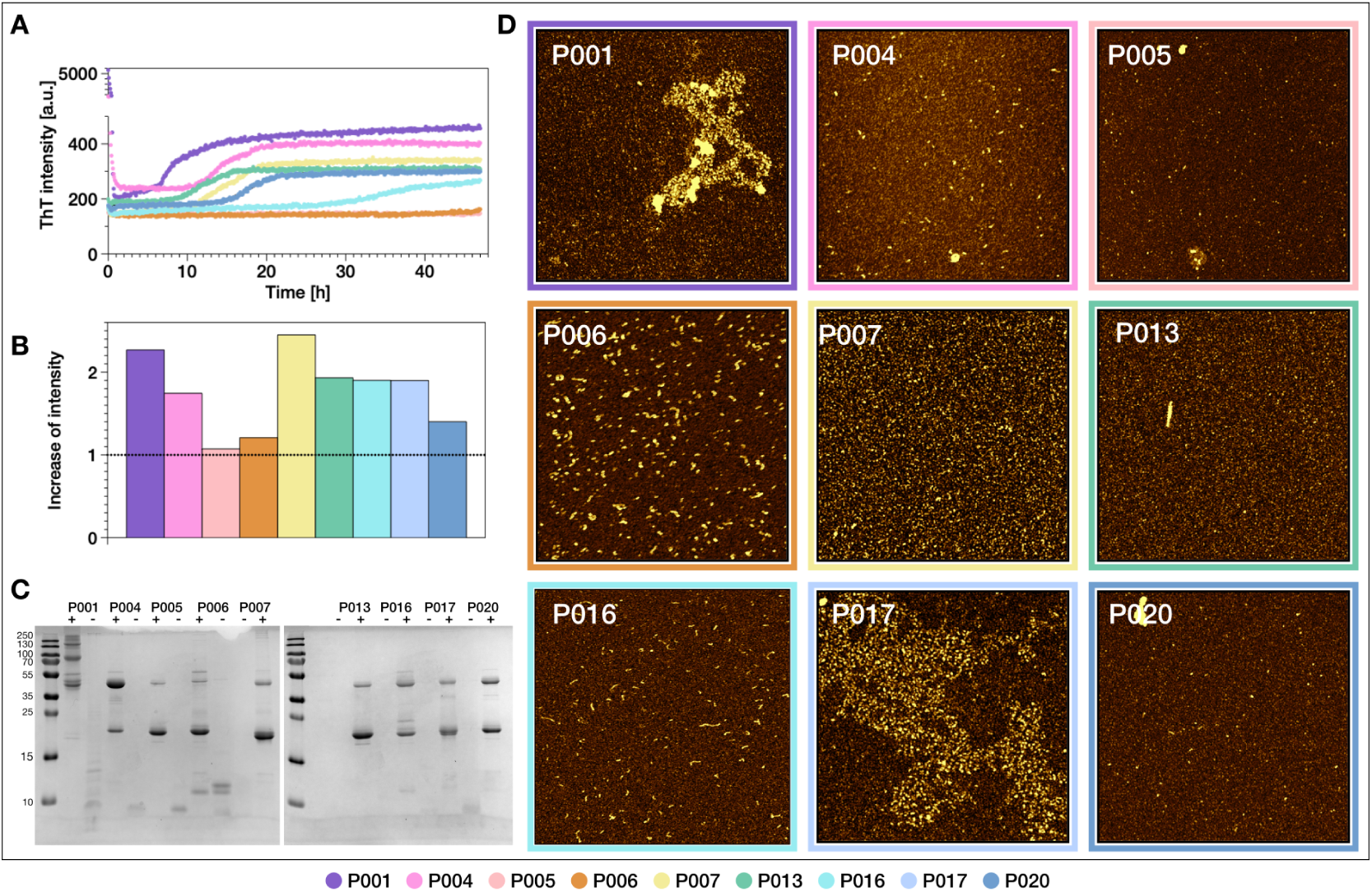
(A) ThT-fluorescence aggregation assay of the IgLC samples at pH 3 in the presence of 10 μM pepstatin A and 10 μM E-64 measured in a high-binding surface plate at 37°C and (B) the extracted factor of the intensity increase. (C) The light chains were analysed by non-reducing SDS-PAGE after incubation for 50 h with inhibitors (+) and without (-). (D) AFM-height-images of the products of the aggregation assay after 50 h. The image scale is 2 × 2 μm. The colour range represents the height from −1 to 5 nm.

The influence of the two different protease inhibitors was further examined at pH 3 and pH 4 for the aggregation of P005 and P016 under agitation conditions (supplementary figure 11). The light chains were incubated with 10 μM or 1 μM of both inhibitors or with 1 μM of only one inhibitor. E-64 alone showed no effect on the aggregation kinetics of P005, but prolonged the aggregation of P016 at pH 3. Even though there was only a weak increase in ThT signal, we could confirm the presence of fibrillar structures in P016 at pH 3 in the presence of the inhibitors. In addition to the fibrils, also some small elongated oligomers could be seen in the AFM images of P016 incubated with inhibitor. Finally, we tested whether the fibrils which are formed in the absence of protease inhibitors can induce the aggregation of the intact light chain. Therefore we incubated the LC proteins at 55°C, where the IgLCs are partly to fully unfolded (see supplementary figure 5) and added seeds which were produced at pH 3 or pH 4 (supplementary figure 12). For P006 and P017, we indeed observed an increase in ThT fluorescence intensity, which was not observed in unseeded samples.

## Discussion

The problem of rationalising the aggregation behaviour of immunoglobulin light chains is a formidable one, given the fact that every single patient will present a light chain with a different amino acid sequence. In particular the question as to which factors determine whether a given light chain forms amyloid fibrils or amorphous aggregates has been extensively studied in recent years. Various biochemical and biophysical properties, such as thermodynamic stability^13,15,20,64,65^, propensity to form dimers^23,25^, ability to refold^15^ or proteolytic digestability as a proxy for protein dynamics^26^ have been proposed to correlate with one or the other type of aggregate formation. In this study, we have performed an in depth characterisation of these and additional factors for 9 patient-derived light chains. A complete overview over the data can be found in table 1. None of these 9 samples stems from an amyloidosis patient, and yet the aforementioned biophysical and biochemical properties span the full range of dimerisation (0.1-1), thermal stability (50-64°C), efficiency of refolding (0-0.7) and trypsin digestability (0.1-1) previously proposed to correlate with amyloid fibril formation in patients. In addition to these parameters, we also characterised the overall aggregation behaviour, induced by heat and low pH, at unprecedented detail. Overall, our ThAgg-Fip approach yields a unique thermodynamic and aggregation fingerprint of each of the IgLCs of our study. While different patient-derived IgLCs may appear similar if characterised only in a single dimension, such as thermal stability or proteolytic digestibility, a multiparametric investigation, such as the one we present here, highlights the uniqueness of each IgLCs. However, despite this uniqueness in overall behaviour, we made the remarkable discovery that every one of the IgLCs of our study can form amyloid fibrils under physiologically relevant conditions. Mildly acidic pH values can be encountered by IgLCs in the kidney due to lysosomal proteolysis^66^ and a decreased pH with a progressive chronic kidney disease (CKD)^67^. Low pH values have also previously been reported to facilitate IgLC aggregation^15,68^. We were able to confirm the formation of amyloid fibrils within all investigated IgLCs, using ThT-fluorescence assays and AFM. Furthermore, we were able to identify the presence of proteases, most notably cathepsins, as decisive for amyloid fibril formation, because of their ability to fragment the full-length protein under physiologically relevant mildly acidic conditions. The finding that all our investigated samples are able to form amyloid fibrils is even more remarkable, as none of our IgLCs appear to form amyloid fibrils in the patient. In order to understand this discrepancy, we determined the amino acid sequences of all 9 samples investigated in this work, and in addition the sequence of the single IgLC from an amyloidosis patient (P011) in our larger data set^19^. The sequences of the IgLCs were determined by a combination of top-down and bottom-up proteomics with specific data analysis for patient-derived light chains. The development and details of this workflow are the subject of a separate publication^41^, where we also discuss some of the specific features of the individual IgLCs. An alignment of the full length amino acid sequences of the two λ light chains P001 and P011 and of the eight *κ* light chains P004, P005, P006, P007, P013, P016, P017 and P020 are presented in the supplementary materials (supplementary figures 1 and 2). The constant regions of the *κ* light chains are identical, apart from P013, which has a valine mutated to a leucine. An alignment of only the variable regions can be found in Figure 7. The sequences contain five cysteine residues which stabilise the monomers by two disulfide bridges between C23 and C87-89 and between C133-C135 and C193-195. The fifth cysteine is cysteinylated in the monomer and is responsible for the disulfide bridge in the dimer. However P001 contains two additional adjacent cysteines at position 99 and 100. Besides the displayed sequence we detected by MS analysis an additional IgLC proteoform in two samples (P004 and P017), and we were able to determine the sequence of the second isoform in P004, which contains two homodimers and one heterodimer, while P017 contains only one homodimer and one heterodimer. By means of the intact mass analysis by mass spectrometry we detected in the sample P011, which stems from the only confirmed amyloidosis patient in our cohort, two other proteins with a similar mass of 20.9 and 21.1 kDa at high abundance in addition to the light chain. This contamination prevented a detailed and accurate biophysical analysis of this sample. Therefore we excluded P011 from the further experiments and analysed solely its amino acid sequence. We used the international immunogenetics information system (IMGT) to search for the germline sequences of the investigated light chains. Both lambda IgLCs have a different origin; P001 from IGLV2-11 and P011 from IGLV1-40, which explains the large differences between the sequences (Figure 7 A), supplementary figure 1). P001 is very similar to the germline sequence; it contains only three mutations: from a serine and an asparagine to a threonine and an additional threonine to alanine mutation. The *κ* IgLCs origin from four different germline sequences. P005 from IGKV1-39, P004 from IGKV3-20, P007, P013 and P020 from IGKV1-5 and P006, P016 and P017 from IGKV1-33 (Figure 7 B). The secondary structure of light chains is dominated by *β*-strands. In order to be able to judge a possible influence of the mutations on the native structures, we searched the PDB for three-dimensional structures of the variable regions of the appropriate germline sequence and highlighted the positions of the mutated amino acids (Figure 7). The sequence changes of P004 and P005 could have a significant impact on the structure, because they affect different *β* strand regions. The sequence changes identified for P013, P020, P016 and P017, on the other hand, are mainly in the loop regions. For some of our sample (P001, P006, P013, P016, P020), we used a top-down proteomics approach to determine the sequence of LC fragments formed at pH 3 and 3. Truncated LC sequences could therefore be determined with high precision and allowed to identify wich parts of the sequences formed amyloid fibrils (supplementary Figure 18). We found different numbers of fragments, and both the constant and variable domains to be represented. The central regions of the IgLCs were almost never found to be part of the fibrils. It is interesting to compare these results with the structures of IgLC amyloid fibrils extracted from patients that have recently been solved by cryo-EM. Fibrils of a λ 1 AL IgLC from an explanted heart were found to consist of a 91-residue fragment of the variable domain^69^, whereas in another fibril structure, a slightly shorter fragment of 77 amino acids of the variable domain was found to form the fibril core^70^. In our fibrils formed *in vitro* we only detected a fragment of almost the entire variable domain in P006 (pH 3), in addition to shorter fragments of the same region. We could find many similar C-terminal fragments, which is in accordance with the heterogeneous fragmentation positions reported recently^71^. The fragments we have detected are mostly rather short with an average molecular weight of 3.8 kDa. Previous studied already demonstrated that short peptides of 12 residues of the variable and constant domain of an amyloidogenic λ LC can form fibrillar structures *in vitro*^72^. It has been proposed that at least some proteolytic cleavage events happen after the assembly into amyloid fibrils *in vivo*^71,73^. This is likely also to be the case for our amyloid fibrils assembled *in vitro*. However, our demonstration that some degree of proteolytic cleavage is required to initiate the formation of amyloid fibrils has also been reported for IgLCs from amyloidosis patients^74,75^, and is therefore not restricted to IgLCs from MM patients. A previous study applied computational predictions using ZipperDB to identify so called steric zippers that drive the assembly of amyloid fibrils in IgLCs^17^. Our determination of the amino acid sequences enabled us to apply various bioinformatic prediction tools with the aim to rationalise the universal ability of our samples to form amyloid fibrils.

**Figure 7.**
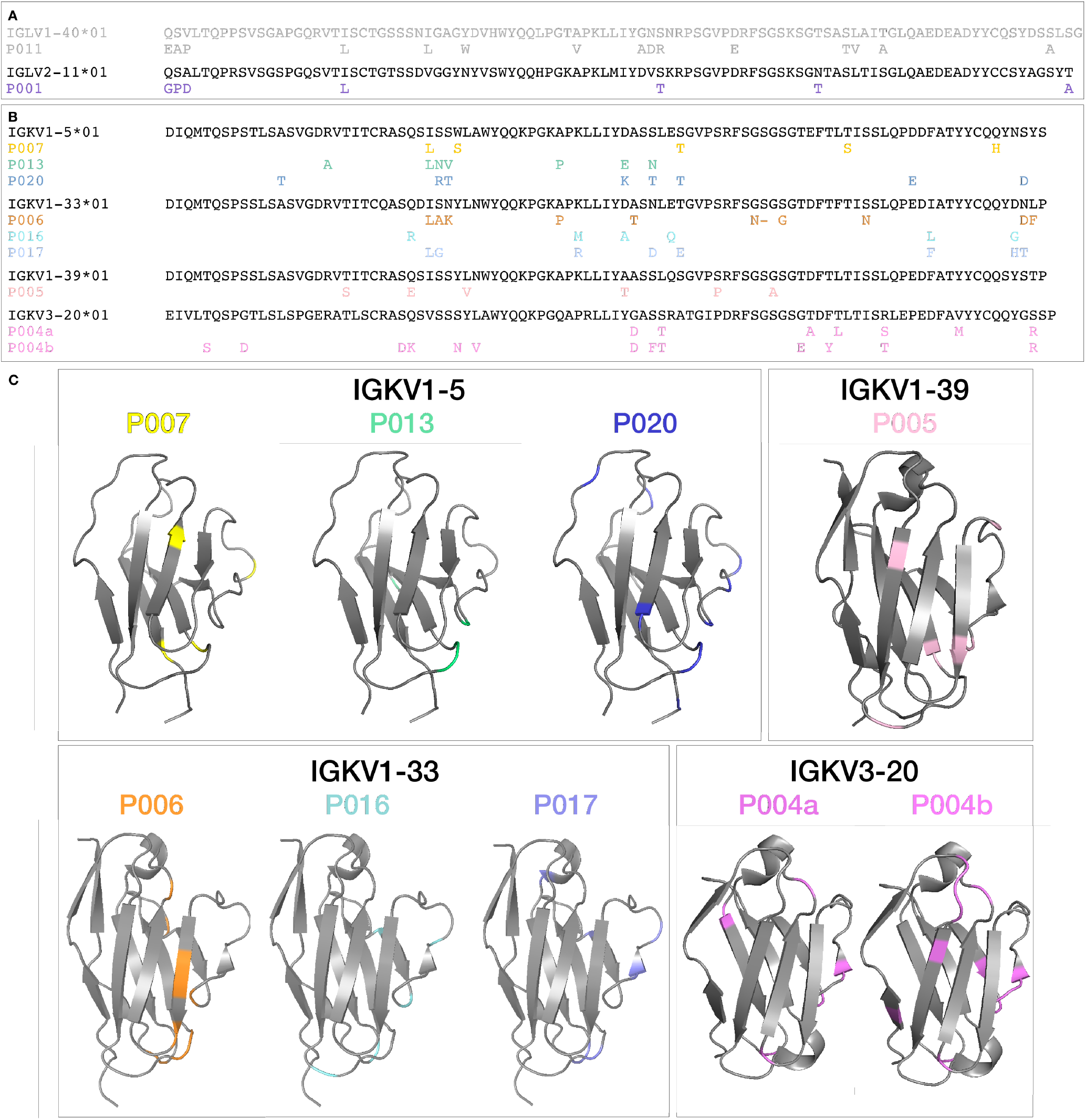
Sequences of the (A) λ and (B) *κ* variable domain of the germline sequences. The mutations in the sequences of the IgLC samples compared to the corresponding germline sequence are labelled. Mutations between isoleucine and leucine are not indicated due to the uncertainty of sequencing. (C) Three-dimensional structure of the variable regions of the *κ* germline sequences from the RSCB protein data bank (5I1C, 3CDF, 3UPA) and UniProtKB (P01602) visualized with PyMOL 2.4. (Schrödinger). The mutated amino acids in the investigated IgLCs are highlighted in colour.

**Figure 8.**
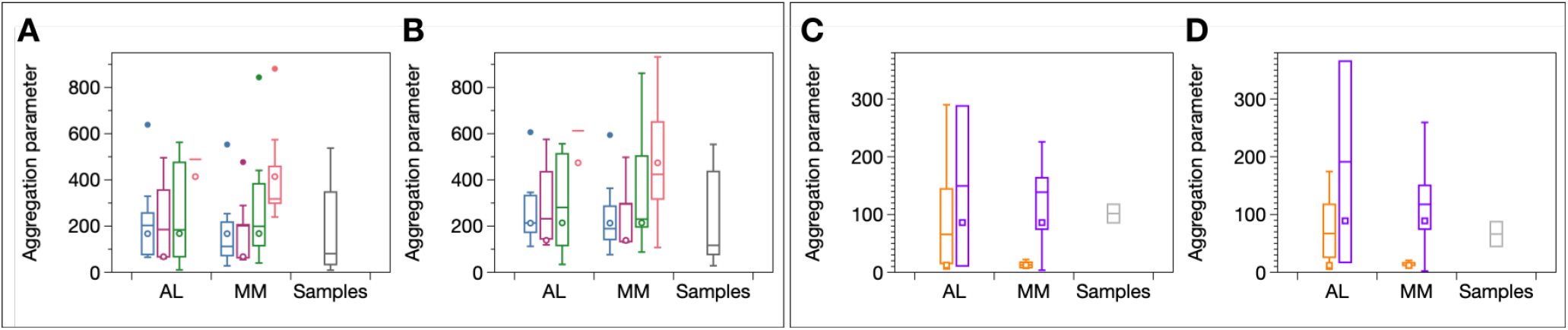
Variable *κ* region sequences were analysed with Tango at pH 7 (A) and pH 3 (B). The sequences, which originate from different germline sequences were selected from AL-Base and were categorised according to whether the patient was suffering from AL-Amyloidosis or MM. IGKV1-5 (blue) AL n=10, MM n=9; IGKV1-33 (violet) AL n=8, MM n=10; IGKV1-39 (green) AL n=10, MM n=10; IGKV3-20 (orange) AL n=7, MM n=10. Variable lambda region sequences were analysed with Tango at pH 7 (C) and pH 3 (D). The sequences, which originate from different germline sequences were selected from AL-Base and were categorised according to whether the patient was suffering from AL-Amyloidosis or MM. IGLV1-40 (orange) AL n=10, MM n=5; IGLV2-11 (violet) AL n=2, MM n=11. The open circles indicate the germline sequence.

We tested the amyloid propensity of the sequences with several freely available computational algorithms such as ZipperDB^76^, Waltz^77^, Tango^78^ and Pasta^79^ (supplementary figures 13-16). Tango revealed the largest differences within the sequences, whereas Pasta displayed no amyloid potential in the variable domains. Furthermore, we also found that the sequence regions identified to be part of the fibrils do not systematically align with the ‘hot spots’ of aggregation propensity and amyloidogenicity predicted by the various algorithms. In addition to our set of sequences, we also determined the aggregation propensities of a range of *κ* and λ sequences from different germlines selected from AL-Base^42^ as a representative collection. The sequences were categorized depending on whether the patient was suffering from AL-Amyloidosis or multiple myeloma. The tested algorithms did not identify any clear distinctions between amyloidogenic and non-amyloid forming light chains.

These results derived from state of the art amyloid prediction algorithms validate our findings that the origin of the differential amyloidogenicity of IgLCs in patients is not primarily determined by their amino acid sequence. Our main discovery can therefore be summarised as follows. All immunoglobulin light chains have a comparable and high intrinsic amyloidgenic potential, and the reason that only some are able to realise this potential is to be found in the complex interplay between intrinsic physical properties of the peptide and environmental conditions *in vivo*. In particular the presence and action of proteases such as cathepsins in a patient facilitates the deposition of the IgLCs in the form of amyloid fibrils. This conclusion corresponds to a major paradigm shift in the field of IgLC aggregation, where the origin of *in vivo* deposition behaviour has almost exclusively been searched for in the sequence alone. Our findings might also hint towards a possible mechanism for the preferential deposition of light chains in specific organs that is often observed. The biochemical micro-environment conducive to IgLC cleavage is likely to differ both between patients, as well as in different organs within a given patient. It is also important to stress that it may not be necessary to cleave the light chain quantitatively in order to allow the conversion into amyloid fibrils. In our *in vitro* experiments, we provided optimal conditions for efficient protolytic cleavage, leading to relatively short sequence fragments (see supplementary figure 18) which are apparently able to form amyloid fibrils particularly readily. We have provided evidence in our study that full length IgLCs can be seeded in some cases with fibrils formed under conditions that favour cleavage (see supplementary figure 12). While this seeding could be very inefficient, it may still be relevant over clinical time scales. We believe that these findings have the potential to have a major impact on how the problem of AL amyloidosis will be tackled in the future. While our results, therefore, represent a major advance in our understanding of AL amyloidosis, the more general question about absolute IgLC solubility in vivo still remains unanswered. Some over-produced IgLCs are very highly soluble and do not cause any damage even if present at gram scale in the patient’s plasma and urine, while others are highly insoluble and can form deposits, which among other things, can cause severe kidney damage. In a previous study we correlated the severity of kidney damage of the patients, with the biophysical and biochemical characteristics of the IgLCs^19^. Using simple pair-wise correlations between a small number of paramters, we were unable to find any clear relationship. In the present, significantly extended study, we employed ThAgg-fingerprinting and find that apart from the universal amyloidogenicity, every IgLC has a unique fingerprint. Our results, which show a large spread of the values of the individual parameters, render it very unlikely that a single biophysical or biochemical parameter can be found to correlate with the complex *in vivo* solubility behaviour of IgLCs. The multidimensional nature of this problem requires a multiparametric approach, such as the ThAgg-Fip method we present here. In supplementary figures 18-20, we show full correlation matrices (Pearson and Spearman) and a scatter matrix between the full set of biophysical and biochemical protein properties and the clinical parameters of the patients. This relatively simple analysis already reveals a plethora of potential links between seemingly disconnected parameters. The potential of this approach is further exemplified by supplementary figure 21, where the clinical GFR-CKD-EPI parameter, a measure for kidney function, is predicted by a linear model using the three parameters trypsin digestibility, ΔG, and dT_*agg*_/dlog *c*, with a Pearson correlation of 0.91. While such preliminary correlations are intriguing, they should only be interpreted with care. The low amount of data points did not allow for independent data to validate the model, which makes it difficult to control for over-fitting. Also, the large number (>10) of largely orthogonal protein properties we have been able to measure for each of our IgLCs renders such an analysis prone to occurrences of Freedman’s paradox^80^ and hence necessitates a sample size at least 5-10 times larger than the one we have been able to assemble for the present study. We are confident, however, that if ThAgg-Fip is applied at such a scale to patient-derived samples and combined with a maximal amount of patient clinical information (disease severity, proteomics data), this type of statistical analysis has the potential to finally unravel the origin and mechanisms of IgLC solubility and associated protein aggregation and deposition diseases.

## Supporting information

Table with peptide fragments

## Acknowledgements

RSH thanks the Manchot foundation for support. AKB and RKN thank the Novo Nordisk Foundation for support (NN-FSA170028392). This work has been supported by EPIC-XS, project number 823839, funded by the Horizon 2020 programme of the European Union, the Institut Pasteur and CNRS.

## References

1. Braak, H. & Braak, E. Neuropathological stageing of Alzheimer-related changes. Acta neuropathologica 82, 239–259 (1991).

2. Knowles, T. P. J., Vendruscolo, M. & Dobson, C. M. The amyloid state and its association with protein misfolding diseases. Nat. reviews. Mol. cell biology 15, 384–96 (2014).

3. Chiti, F. & Dobson, C. M. Protein misfolding, amyloid formation, and human disease: A summary of progress over the last decade. Annu. Rev. Biochem. 86, 27–68 (2017).

4. Wechalekar, A. D., Gillmore, J. D. & Hawkins, P. N. Systemic amyloidosis. The Lancet 387, 2641–2654 (2016).

5. Bakkus, M., Heirman, C., Van Riet, I., Van Camp, B. & Thielemans, K. Evidence that multiple myeloma Ig heavy chain VDJ genes contain somatic mutations but show no intraclonal variation. Blood 80, 2326–2335 (1992).

6. Chapman, M. A. et al. Initial genome sequencing and analysis of multiple myeloma. Nature 471, 467–472 (2011).

7. Delman, G. & Gally, J. The nature of Bence-Jones proteins: chemical similarities to polypeptide chains of myeloma globulins and normal γ-globulins. The J. experimental medicine 116, 207–227 (1962).

8. Magrangeas, F. et al. Light-chain only multiple myeloma is due to the absence of functional (productive) rearrangement of the IgH gene at the DNA level. Blood 103, 3869–3875 (2004).

9. Herrera, G. A. Renal lesions associated with plasma cell dyscrasias: practical approach to diagnosis, new concepts, and challenges. Arch. pathology & laboratory medicine 133, 249–267 (2009).

10. Sanders, P. W. & Booker, B. B. Pathobiology of cast nephropathy from human Bence Jones proteins. The J. clinical investigation 89, 630–639 (1992).

11. Buxbaum, J. & Gallo, G. Nonamyloidotic monoclonal immunoglobulin deposition disease: light-chain, heavy-chain, and light-and heavy-chain deposition diseases. Hematol. clinics North Am. 13, 1235–1248 (1999).

12. Glenner, G. G., Ein, D. & Terry, W. D. The immunoglobulin origin of amyloid. Am. J. Med. 52, 141–147 (1972).

13. Kim, Y.-s. et al. Thermodynamic modulation of light chain amyloid fibril formation. J. Biol. Chem. 275, 1570–1574 (2000).

14. Arosio, P. et al. In vitro aggregation behavior of a non-amyloidogenic λ light chain dimer deriving from U266 multiple myeloma cells. PloS one 7 (2012).

15. Blancas-Mejía, L. M. et al. Thermodynamic and fibril formation studies of full length immunoglobulin light chain AL-09 and its germline protein using scan rate dependent thermal unfolding. Biophys. Chem. 207, 13–20 (2015).

16. Andrich, K. et al. Aggregation of full-length immunoglobulin light chains from systemic light chain amyloidosis (AL) patients is remodeled by epigallocatechin-3-gallate. J. Biol. Chem. 292, 2328–2344 (2017).

17. Brumshtein, B. et al. Identification of two principal amyloid-driving segments in variable domains of Ig light chains in systemic light-chain amyloidosis. J. Biol. Chem. 293, 19659–19671 (2018).

18. Weber, B. et al. The antibody light-chain linker regulates domain orientation and amyloidogenicity. J. Mol. Biol. 430, 4925–4940 (2018).

19. Sternke-Hoffmann, R. et al. Biochemical and biophysical characterisation of immunoglobulin free light chains derived from an initially unbiased population of patients with light chain disease. PeerJ 8, e8771 (2020).

20. Raffen, R. et al. Physicochemical consequences of amino acid variations that contribute to fibril formation by immunoglobulin light chains. Protein Sci. 8, 509–517 (1999).

21. Klimtchuk, E. S. et al. The critical role of the constant region in thermal stability and aggregation of amyloidogenic immunoglobulin light chain. Biochemistry 49, 9848–9857 (2010).

22. González-Andrade, M. et al. Mutational and genetic determinants of λ6 light chain amyloidogenesis. FEBS J. 280, 6173–6183 (2013).

23. Gatt, M. E. et al. The use of serum free light chain dimerization patterns assist in the diagnosis of al amyloidosis. Br. J. Haematol. 182, 86–92 (2018).

24. Qin, Z., Hu, D., Zhu, M. & Fink, A. L. Structural characterization of the partially folded intermediates of an immunoglobulin light chain leading to amyloid fibrillation and amorphous aggregation. Biochemistry 46, 3521–3531 (2007).

25. Bernier, G. M. & Putnam, F. W. Monomer–dimer forms of Bence Jones proteins. Nature 200, 223–225 (1963).

26. Oberti, L. et al. Concurrent structural and biophysical traits link with immunoglobulin light chains amyloid propensity. Sci. Rep. 7, 1–11 (2017).

27. Mukherjee, S., Pondaven, S. P., Hand, K., Madine, J. & Jaroniec, C. P. Effect of amino acid mutations on the conformational dynamics of amyloidogenic immunoglobulin light-chains: A combined NMR and in silico study. Sci. Rep. 7, 1–13 (2017).

28. Kazman, P. et al. Fatal amyloid formation in a patient’s antibody light chain is caused by a single point mutation. Elife 9, e52300 (2020).

29. Sakano, H., Hüppi, K., Heinrich, G. & Tonegawa, S. Sequences at the somatic recombination sites of immunoglobulin light-chain genes. Nature 280, 288–294 (1979).

30. Lefranc, M.-P. & Lefranc, G. The immunoglobulin factsbook (Academic press, 2001).

31. Teng, G. & Papavasiliou, F. N. Immunoglobulin somatic hypermutation. Annu. Rev. Genet. 41, 107–120 (2007).

32. Gearhart, P. J., Johnson, N. D., Douglas, R. & Hood, L. IgG antibodies to phosphorylcholine exhibit more diversity than their IgM counterparts. Nature 291, 29–34 (1981).

33. McBride, O. W. et al. Chromosomal location of human kappa and lambda immunoglobulin light chain constant region genes. J. Exp. MEd. 155, 1480–1490 (1982).

34. Malcolm, S. et al. Localization of human immunoglobulin kappa light chain variable region genes to the short arm of chromosome 2 by in situ hybridization. Proc. Natl. Acad. Sci. 79, 4957–4961 (1982).

35. Isobe, T. & Osserman, E. F. Patterns of amyloidosis and their association with plasma-cell dyscrasia, monoclonal immunoglobulins and Bence-Jones proteins. New Engl. J. Med. 290, 473–477 (1974).

36. Scaviner, D., Barbié, V., Ruiz, M. & Lefranc, M.-P. Protein displays of the human immunoglobulin heavy, kappa and lambda variable and joining regions. Exp. Clin. Immunogen. 16, 234–240 (1999).

37. Upadhyay, A. A. et al. BALDR: a computational pipeline for paired heavy and light chain immunoglobulin reconstruction in single-cell RNA-seq data. Genome medicine 10, 1–18 (2018).

38. Srzentic, K. et al. Interlaboratory study for characterizing monoclonal antibodies by top-down and middle-down mass spectrometry. J. Am. Soc. for Mass Spectrom. 31, 1783–1802 (2020).

39. Tran, N. H. et al. Complete de novo assembly of monoclonal antibody sequences. Sci. Rep. 6, 1–10 (2016).

40. Dupré, M. et al. Optimization of a top-down proteomics platform for closely related pathogenic bacterial discrimination. J. Proteome Res. (2020).

41. Dupré, M. et al. De novo sequencing of antibody light chain proteoforms from patients with multiple myeloma, 10.26434/chemrxiv.14541711.v1 (2021).

42. Bodi, K. et al. AL-base: a visual platform analysis tool for the study of amyloidogenic immunoglobulin light chain sequences. Amyloid 16, 1–8 (2009).

43. Schuck, P. Size-distribution analysis of macromolecules by sedimentation velocity ultracentrifugation and lamm equation modeling. Biophys. journal 78, 1606–1619 (2000).

44. Brown, P. H. & Schuck, P. Macromolecular size-and-shape distributions by sedimentation velocity analytical ultracentrifugation. Biophys. J. 90, 4651–4661 (2006).

45. Putnam, F. W., Easley, C. W., Lynn, L. T., Ritchie, A. E. & Phelps, R. A. The heat precipitation of Bence-Jones proteins. I. optimum conditions. Arch. Biochem. Biophys. 83, 115–13 (1959).

46. Žoldák, G., Jancura, D. & Sedlák, E. The fluorescence intensities ratio is not a reliable parameter for evaluation of protein unfolding transitions. Protein Sci. 26, 1236–1239 (2017).

47. Hamborg, L. et al. Global analysis of protein stability by temperature and chemical denaturation. Anal. Biochem. 605, 113863 (2020).

48. Lindorff-Larsen, K. Dissecting the statistical properties of the linear extrapolation method of determining protein stability. Protein Eng. Des. Sel. 32, 471–479 (2019).

49. Nielsen, L. et al. Effect of environmental factors on the kinetics of insulin fibril formation: elucidation of the molecular mechanism. Biochemistry 40, 6036–6046 (2001).

50. Arosio, P. et al. Microfluidic diffusion analysis of the sizes and interactions of proteins under native solution conditions. ACS nano 10, 333–341 (2015).

51. Yates, E. V. et al. Latent analysis of unmodified biomolecules and their complexes in solution with attomole detection sensitivity. Nat. Chem. 7, 802–809 (2015).

52. Kunz, P. et al. The structural basis of nanobody unfolding reversibility and thermoresistance. Sci. Reports 8, 7934 (2018).

53. Stagg, L., Zhang, S.-Q., Cheung, M. S., WBence Jones proteinittung-Stafshede, P. & Molecular crowding enhances native structureand stability of α/β protein flavodoxin. Proc. Nat. Ac. Sc. 104, 18976–18981 (2007).

54. Mohamed, M. M. & Sloane, B. F. Multifunctional enzymes in cancer. Nat. Rev. Cancer 6, 764–775 (2006).

55. Turk, V. et al. Cysteine cathepsins: from structure, function and regulation to new frontiers. Biochimica et Biophys. Acta (BBA)-Proteins Proteomics 1824, 68–88 (2012).

56. Giusti, I. et al. Cathepsin B mediates the pH-dependent proinvasive activity of tumor-shed microvesicles. Neoplasia 10, 481–488 (2008).

57. Zhao, B. et al. Crystal structure of human osteoclast cathepsin K complex with E-64. Nat. Struct. Biol. 4, 109–111 (1997).

58. Hashida, S., Towatari, T., Kominami, E. & Katunuma, N. Inhibitions by E-64 derivatives of rat liver cathepsin B and cathepsin L in vitro and in vivo. J. Biochem. 88, 1805–1811 (1980).

59. Schaschke, N., Assfalg-Machleidt, I., Machleidt, W., Turk, D. & Moroder, L. E-64 analogues as inhibitors of cathepsin B. on the role of the absolute configuration of the epoxysuccinyl group. Bioorg. Med. Chem. 5, 1789–1797 (1997).

60. Barrett, A. J. et al. L-trans-Epoxysuccinyl-leucylamido (4-guanidino) butane (E-64) and its analogues as inhibitors of cysteine proteinases including cathepsins B, H and L. Biochem. J. 201, 189–198 (1982).

61. Santamaría, I., Velasco, G., Pendás, A. M., Fueyo, A. & López-Otín, C. Cathepsin Z, a novel human cysteine proteinase with a short propeptide domain and a unique chromosomal location. J. Biol. Chem. 273, 16816–16823 (1998).

62. Whitaker, J., Herman, P., Sparacio, S., Zhou, S. & Benveniste, E. Changes induced in astrocyte cathepsin D by cytokines and leupeptin. J. Neurochem. 57, 406–414 (1991).

63. Chen, C.-S., Chen, W.-N. U., Zhou, M., Arttamangkul, S. & Haugland, R. P. Probing the cathepsin D using a BODIPY FL-pepstatin A: applications in fluorescence polarization and microscopy. J. Biochem. Biophys. Meth. 42, 137–151 (2000).

64. Hernández-Santoyo, A. et al. A single mutation at the sheet switch region results in conformational changes favoring λ6 light-chain fibrillogenesis. J. Mol. Biol. 396, 280–292 (2010).

65. Nowak, M. Immunoglobulin kappa light chain and its amyloidogenic mutants: a molecular dynamics study. Proteins: Struct. Funct. Bioinforma. 55, 11–21 (2004).

66. Pillay, C. S., Elliott, E. & Dennison, C. Endolysosomal proteolysis and its regulation. Biochem. J. 363, 417–429 (2002).

67. Nakanishi, N. et al. Low urine pH is a predictor of chronic kidney disease. Kidney Blood Press. Res 35, 77–81 (2012).

68. Rostagno, A. et al. pH-dependent fibrillogenesis of a V*κ*III Bence Jones protein. Brit. J. Haematol. 107, 835–843 (1999).

69. Radamaker, L. et al. Cryo-EM structure of a light chain-derived amyloid fibril from a patient with systemic AL amyloidosis. Nat. communications 10, 1–8 (2019).

70. Swuec, P. et al. Cryo-em structure of cardiac amyloid fibrils from an immunoglobulin light chain al amyloidosis patient. Nat. communications 10, 1269 (2019).

71. Lavatelli, F. et al. Mass spectrometry characterization of light chain fragmentation sites in cardiac AL amyloidosis: insights into the timing of proteolysis. J. Biol. Chem. 295, 16572–16584 (2020).

72. Schmidt, A., Annamalai, K., Schmidt, M., Grigorieff, N. & Fändrich, M. Cryo-EM reveals the steric zipper structure of a light chain-derived amyloid fibril. Proc. Natl. Acad. Sci. 113, 6200–6205 (2016).

73. Enqvist, S., Sletten, K. & Westermark, P. Fibril protein fragmentation pattern in systemic AL-amyloidosis. The J. Pathol. A J. Pathol. Soc. Gt. Br. Irel. 219, 473–480 (2009).

74. Eulitz, M., Breuer, M. & Linke, R. P. Is the formation of AL-type amyloid promoted by structural peculiarities of immunoglobulin L-chains? Primary structure of an amyloidogenic λ-L-chain (BJP-ZIM). Biol. Chem. 368, 863–870 (1987).

75. Stevens, F. J. et al. A molecular model for self-assembly of amyloid fibrils: immunoglobulin light chains. Biochem. 34, 10697–10702 (1995).

76. Goldschmidt, L., Teng, P. K., Riek, R. & Eisenberg, D. Identifying the amylome, proteins capable of forming amyloid-like fibrils. Proc. Nat. Acad. Sc. 107, 3487–349 (2010).

77. Maurer-Stroh, S. et al. Exploring the sequence determinants of amyloid structure using position-specific scoring matrices. Nat. Methods 7, 237–242 (2010).

78. Fernandez-Escamilla, A.-M., Rousseau, F., Schymkowitz, J. & Serrano, L. Prediction of sequence-dependent and mutational effects on the aggregation of peptides and proteins. Nat. Biotech. 22, 1302–1306 (2004).

79. Walsh, I., Seno, F., Tosatto, S. C. & Trovato, A. PASTA 2.0: an improved server for protein aggregation prediction. Nucl. Ac. Res. 42, W301–W307 (2014).

80. Freedman, D. A. A note on screening regression equations. The Am. Stat. 37, 152–155 (1983).

