## Supplementary material for "Universal amyloidogenicity of patient-derived immunoglobulin light chains": Table with peptide fragments

| Found in Sample | Sequence | # PrSMs | Theo. Mass [Da] | -Log P-Score | % Residue Cleavages | DeltaMass [ppm] |
|---|---|---|---|---|---|---|
| P001_pH3 | LSLTPEQWKSHRSYSCQVTHEGSTVEKTVA | 1 | 3,445 | 47.44 | 55.2 | 4.06 |
| P001_pH3 | LSLTPEQWKSHRSYSCQVTHEGSTVE | 1 | 3,045 | 41.64 | 72 | 3.79 |
| P001_pH3 | YDVTKRPSGVPDRFSGSKSGTTAS | 1 | 2,499 | 52.93 | 73.9 | 5.57 |
| P001_pH3 | TVLGQPKAAPSVTLFPPSS | 1 | 1,896 | 50.13 | 83.3 | 4.24 |
| P001_pH3 | YYCCSYAGLDLF | 1 | 1,531 | 33.91 | 81.8 | 3.9 |
| P001_pH3 | GPDLTQPRSVSGSPG | 1 | 1,454 | 13.46 | 42.9 | 4.39 |
| P001_pH3 | VCLLSDFYPQVT | 1 | 1,441 | 20.27 | 63.6 | 3.89 |
| P006_pH3 | LQMTQSPSSLSASVGDRVTLTCQASQDLAKYLNWYQQKPGKPPKLLLYDTSNLETGVPSRFSNGGGTDFTFTLNSLQPEDLATYYCQQYDDFPLTFGPGTKVDLKRTVAAPSVF | 1 | 12,643 | 13.34 | 11.5 | 2.35 |
| P006_pH3 | ETGVPSRFSNGGGTDFTFTLNSLQPEDLATYYCQQYDDFPLTFGPGTKVDLKRTVAAPSVF | 1 | 6,722 | 58.53 | 48.3 | 1.68 |
| P006_pH3 | ETGVPSRFSNGGGTDFTFTLNSLQPEDLATYYCQQYDDFPLTFGPGTKVDLKRTVAA | 1 | 6,292 | 17.26 | 17.9 | 4.38 |
| P006_pH3 | DLQMTQSPSSLSASVGDRVTLTCQASQDLAKYLNWYQQKPGKPPKLLL | 2 | 5,361 | 93.47 | 61.7 | -0.77 |
| P006_pH3 | DLQMTQSPSSLSASVGDRVTLTCQASQDLAKYLNWYQQKPGKPPKL | 2 | 5,135 | 80.56 | 64.4 | 4.77 |
| P006_pH3 | SASVGDRVTLTCQASQDLAKYLNWYQQKPGKPPKLLLYDTSN | 1 | 4,753 | 44.52 | 43.9 | 3.33 |
| P006_pH3 | TQSPSSLSASVGDRVTLTCQASQDLAKYLNWYQQKPGKPPKL | 1 | 4,647 | 76.45 | 51.2 | 3.85 |
| P006_pH3 | SASVGDRVTLTCQASQDLAKYLNWYQQKPGKPPKL | 3 | 3,947 | 119.05 | 85.3 | 4.5 |
| P006_pH3 | AKVQWKVDNALQSGNSQESVTEQDSKDSTYSLSSTL | 1 | 3,930 | 82.51 | 71.4 | 6.35 |
| P006_pH3 | KVQWKVDNALQSGNSQESVTEQDSKDSTYSLS | 1 | 3,558 | 12.67 | 16.1 | 3.63 |
| P006_pH3 | AKVQWKVDNALQSGNSQESVTEQDSKDSTYSL | 8 | 3,542 | 112.51 | 80.6 | 3.91 |
| P006_pH3 | LATYYCQQYDDFPLTFGPGTKVDLKRTVAA | 1 | 3,438 | 32.75 | 51.7 | 3.44 |
| P006_pH3 | YYCQQYDDFPLTFGPGTKVDLKR | 1 | 2,810 | 49.31 | 72.7 | 4.48 |
| P006_pH3 | QQYDDFPLTFGPGTKVDLKRTVAA | 1 | 2,666 | 82.58 | 91.3 | 4.68 |
| P006_pH3 | SKADYEKHKVYACEVTHQGLSS | 1 | 2,536 | 64.18 | 90.5 | 4.72 |
| P006_pH4 | VVCLLNNFYPREAKVQWKVDNALQSGNSQESVTEQDSKDSTYSLSSTLTL | 1 | 5,649 | 30.89 | 32.7 | 4.87 |
| P006_pH4 | LNNFYPREAKVQWKVDNALQSGNSQESVTEQDSKDSTYSLSSTLTL | 2 | 5,178 | 16.16 | 26.7 | 8.11 |
| P006_pH4 | LNNFYPREAKVQWKVDNALQSGNSQESVTEQDSKDSTYSLSSTL | 1 | 4,963 | 30.22 | 44.2 | 4.17 |
| P006_pH4 | LNNFYPREAKVQWKVDNALQSGNSQESVTEQDSKDSTYSL | 1 | 4,575 | 19.02 | 28.2 | 4.57 |
| P006_pH4 | AKVQWKVDNALQSGNSQESVTEQDSKDSTYSLSSTLTL | 2 | 4,144 | 30.87 | 43.2 | 5.09 |
| P006_pH4 | ALQSGNSQESVTEQDSKDSTYSLSSTLTL | 1 | 3,075 | 61.64 | 82.1 | 3.29 |
| P013_pH3 | SVVCLLNNFYPREAKVQWKVDNALQSGNSQESVTEQDSKDSTYSLSSTL | 1 | 5,522 | 63.76 | 52.1 | 5.69 |
| P013_pH3 | VVCLLNNFYPREAKVQWKVDNALQSGNSQESVTEQDSKDSTYSLSSTL | 1 | 5,435 | 102.9 | 70.2 | 1.91 |
| P013_pH3 | SVVCLLNNFYPREAKVQWKVDNALQSGNSQESVTEQDSKDSTYSL | 1 | 5,133 | 135.28 | 81.8 | 3.9 |
| P013_pH3 | VVCLLNNFYPREAKVQWKVDNALQSGNSQESVTEQDSKDSTYSL | 1 | 5,046 | 140.63 | 83.7 | 3.94 |
| P013_pH3 | LNNFYPREAKVQWKVDNALQSGNSQESVTEQDSKDSTYSL | 1 | 4,575 | 141.49 | 89.7 | 3.89 |
| P013_pH3 | AKVQWKVDNALQSGNSQESVTEQDSKDSTYSLSSTL | 1 | 3,930 | 42.1 | 54.3 | 0.9 |
| P013_pH3 | SASVGDAVTLTCRASQSLNVWLAWYQQKPGKPPKL | 1 | 3,856 | 88.06 | 82.4 | 3.11 |
| P013_pH3 | AKVQWKVDNALQSGNSQESVTEQDSKDSTYSLSS | 1 | 3,716 | 10.62 | 24.2 | -7.65 |
| P013_pH3 | VWLAWYQQKPGKPPKLLLYEASNL | 1 | 2,842 | 63.49 | 78.3 | 2.48 |
| P013_pH3 | TLSKADYEKHKLYACEVTHQGLSS | 1 | 2,764 | 20.98 | 43.5 | 4.62 |
| P013_pH3 | YACEVTHQGLSSPVTKSFNRGEC | 1 | 2,626 | 17.89 | 36.4 | 3.99 |
| P013_pH3 | SGSGSGTEFTLTLSSLQPDDF | 1 | 2,145 | 43.75 | 65 | 2.95 |
| P013_pH3 | SVVCLLNNFYPRE | 1 | 1,610 | 12.31 | 41.7 | 3.56 |
| P013_pH3 | LSKADYEKHKL | 1 | 1,331 | 17.48 | 60 | 4.81 |
| P013_pH3 | DLQMTQSPSTL | 1 | 1,220 | 34.64 | 100 | 3.07 |
| P013_pH4 | DLQMTQSPSTLSASVGDAVTLTCRASQSLNVWLAWYQQKPGKPPKL | 1 | 5,058 | 19.35 | 26.7 | 4.31 |
| P013_pH4 | VVCLLNNFYPREAKVQWKVDNALQSGNSQESVTEQDSKDSTYSL | 1 | 5,046 | 76.84 | 69.8 | 5.46 |
| P013_pH4 | LNNFYPREAKVQWKVDNALQSGNSQESVTEQDSKDSTYSL | 1 | 4,575 | 16.21 | 17.9 | 5.2 |
| P016_pH3 | LFPPSDEQLKSGTASVVCLLNNFYPREAKVQWKVDNALQSGNSQESVTEQDSKDSTYSLSSTLTL | 1 | 7,207 | 33.55 | 29.7 | 2.5 |
| P016_pH3 | LFPPSDEQLKSGTASVVCLLNNFYPREAKVQWKVDNALQSGNSQESVTEQDSKDSTYSL | 1 | 6,604 | 73.11 | 58.6 | 2.97 |
| P016_pH3 | LFPPSDEQLKSGTASVVCLLNNFYPREAKVQWKVDN | 1 | 4,149 | 129.26 | 94.3 | 2.37 |
| P016_pH3 | AKVQWKVDNALQSGNSQESVTEQDSKDSTYSLSSTLTL | 1 | 4,144 | 55.19 | 51.4 | 1.93 |
| P016_pH3 | SVVCLLNNFYPREAKVQWKVDN | 1 | 2,678 | 74.1 | 85.7 | 2.58 |
| P016_pH4 | TLSSLQPEDLATYYCQQYGNLPLTFGGGTKVELKGTVAA | 1 | 4,190 | 58.31 | 55.3 | 6.16 |
| P016_pH4 | SASVGDRVTLTCQASRDLSNYLNWYQQKPGKAPML | 1 | 3,954 | 81.16 | 79.4 | 2.54 |
| P016_pH4 | KVQWKVDNALQSGNSQESVTEQDSKDSTYSLS | 1 | 3,558 | 15.31 | 16.1 | 2.73 |
| P016_pH4 | AKVQWKVDNALQSGNSQESVTEQDSKDSTYSL | 4 | 3,542 | 81.32 | 64.5 | 5.01 |
| P016_pH4 | YYCQQYGNLPLTFGGGTKVELKGTVAA | 1 | 2,934 | 14.28 | 34.6 | 3.86 |
| P016_pH4 | SASVGDRVTLTCQASRDLSNYLNW | 1 | 2,712 | 47.14 | 65.2 | 5.06 |
| P016_pH4 | QASRDLSNYLNWYQQKPGKAPML | 1 | 2,707 | 21.02 | 45.5 | -2.24 |
| P016_pH4 | ASRDLSNYLNWYQQKPGKAPML | 1 | 2,579 | 71.16 | 81 | 4.16 |
| P016_pH4 | SRDLSNYLNWYQQKPGKAPML | 1 | 2,508 | 64.61 | 80 | 4.72 |
| P016_pH4 | SASVGDRVTLTCQASRDLSNYL | 1 | 2,412 | 71.27 | 81 | 4.01 |
| P016_pH4 | YGNLPLTFGGGTKVELKGTVAA | 3 | 2,192 | 41.57 | 61.9 | 3.91 |
| P020_pH3 | SVVCLLNNFYPREAKVQWKVDNALQSGNSQESVTEQDSKDSTYSLSSTL | 1 | 5,522 | 60.23 | 43.8 | 3.01 |
| P020_pH3 | SVVCLLNNFYPREAKVQWKVDNALQSGNSQESVTEQDSKDSTYSL | 1 | 5,133 | 12.76 | 20.5 | 4.43 |
| P020_pH3 | AKVQWKVDNALQSGNSQESVTEQDSKDSTYSLSSTLTL | 3 | 4,144 | 43.09 | 45.9 | 4.61 |
| P020_pH3 | AKVQWKVDNALQSGNSQESVTEQDSKDSTYSLSSTL | 6 | 3,930 | 85.52 | 82.9 | 4.08 |
| P020_pH3 | AKVQWKVDNALQSGNSQESVTEQDSKDSTYSLSST | 1 | 3,817 | 71.37 | 64.7 | 5.79 |
| P020_pH3 | AKVQWKVDNALQSGNSQESVTEQDSKDSTYSLSS | 2 | 3,716 | 107.81 | 84.8 | 4.81 |
| P020_pH3 | DLQMTQSPSTLSTSVGDRVTLTCRASQSLRTWL | 1 | 3,695 | 46.77 | 50 | 5.09 |
| P020_pH3 | AKVQWKVDNALQSGNSQESVTEQDSKDSTYSL | 7 | 3,542 | 103.66 | 90.3 | 5.21 |
| P020_pH3 | MTQSPSTLSTSVGDRVTLTCRASQSLRTWL | 1 | 3,339 | 67.52 | 82.8 | 1.97 |
| P020_pH3 | TQSPSTLSTSVGDRVTLTCRASQSLRTWL | 1 | 3,208 | 89 | 96.4 | 4.67 |
| P020_pH3 | STSVGDRVTLTCRASQS | 1 | 1,824 | 56.3 | 81.3 | 5.28 |
| P020_pH4 | LNNFYPREAKVQWKVDNALQSGNSQESVTEQDSKDSTYSLSSTLTL | 1 | 5,178 | 42.01 | 46.7 | 2.43 |
| P020_pH4 | LNNFYPREAKVQWKVDNALQSGNSQESVTEQDSKDSTYSLSSTL | 1 | 4,963 | 27.15 | 39.5 | 6.47 |
| P020_pH4 | LFPPSDEQLKSGTASVVCLLNNFYPREAKVQWKVDN | 1 | 4,149 | 99.65 | 88.6 | 3.82 |
| P020_pH4 | AKVQWKVDNALQSGNSQESVTEQDSKDSTYSLSSTLTL | 3 | 4,144 | 21.12 | 29.7 | 6.73 |
| P020_pH4 | STSVGDRVTLTCRASQSLRTWLAWYQQKPGKA | 1 | 3,651 | 53.38 | 61.3 | 5.02 |
| P020_pH4 | ALQSGNSQESVTEQDSKDSTYSLSSTLTL | 4 | 3,075 | 81.65 | 89.3 | 3 |
| P020_pH4 | STSVGDRVTLTCRASQSLRTWL | 1 | 2,493 | 22.33 | 61.9 | 2.89 |
| P020_pH4 | STSVGDRVTLTCRASQS | 1 | 1,824 | 11.47 | 37.5 | 3.96 |
