## Supplementary Information for "Universal amyloidogenicity of patient-derived immunoglobulin light chains"

### Universal amyloidogenicity of patient-derived immunoglobulin light chains - Supplementary Information

[1]Institut für Physikalische Biologie, Heinrich-Heine Universität Düsseldorf, Düsseldorf, Germany
[2]Forschungszentrum Jülich GmbH, IBI-7, Jülich, Germany
[3]Department of Biotechnology and Biomedicine, Technical University of Denmark, Lyngby, Denmark
[4]Condensed Matter Physics Laboratory, Heinrich-Heine Universität Düsseldorf, Düsseldorf, Germany
[5]Mass Spectrometry for Biology Unit, CNRS USR2000, Institut Pasteur, Paris 75015, France
[6]Cologne Biocenter, Cluster of Excellence on Plant Sciences, Mass Spectrometry Platform, University of Cologne, Cologne, Germany
[7]Department of Hematology, Oncology and Clinical Oncology, Heinrich-Heine Universität Düsseldorf, Düsseldorf, Germany
[8]Forschungszentrum Jülich GmbH, IBI-4, Jülich, Germany

**Table 1.** Patient characteristics at the time of examination. The samples were taken at the time point of diagnosis. The patients are categorised according to their CKD stage: Group I "good" stage 1+2, n=5 (P004, P007, P011, P017, P020), Group II intermediate stage 3, n=4 (P001, P005, P013, P016) and Group III bad stage 4+5, n=1 (P006). Apart from P007, which presented an acute kidney injury, all patients had a chronically damaged kidney.

| | Age | Gender | Disease type | Isotype | FLC Serum [mg/l] | FLC Urine [mg/l] | TPU [g/24 h] | Kidney function creatinin [mg/dl] | GFR-CKD-EPI [ml/min] | CKD Stadium | Kidney function recovered? (Improvement = 30%; yes/no) | Kidney function recovered? Creatinine [mg/dl] |
|---|---|---|---|---|---|---|---|---|---|---|---|---|
| P001 | 47 | ♂ | MM | $\lambda$ | 3750 | 1060 | 13 | 1.3 | 50 | 3 | Yes | 0,9 |
| P004 | 72 | ♂ | MM | $\kappa$ | 5280 | 7650 | 4 | 1.1 | 90 | 1 | Yes | 0,9 |
| P005 | 65 | ♀ | MM | $\kappa$ | 1250 | 6140 | 3 | 1.2 | 48 | 3 | NA | acute kidney injury |
| P006 | 66 | ♀ | MM | $\kappa$ | 2460 | 6880 | 3 | 1.9 | 27 | 4 | Yes | 1,3 |
| P007 | 54 | ♂ | MM | $\kappa$ | 898 | NA | 1 | 1 | 83 | 2 | No | 1,3 |
| P011 | 45 | ♀ | AL | $\lambda$ | 120 | NA | 6.2 | 0.7 | 108 | 1 | No | 0,8 |
| P013 | 65 | ♂ | MM | $\kappa$ | 11000 | NA | 3 | 1.5 | 50 | 3 | No | 1,7 |
| P016 | 59 | ♂ | MM | $\kappa$ | 1150 | 175 | 3 | 1.6 | 45 | 3 | No | 1,23 |
| P017 | 72 | ♀ | MM | $\kappa$ | 1380 | NA | 4 | 0.67 | 90 | 1 | No | 0,8 |
| P020 | 64 | ♀ | MM | $\kappa$ | 4120 | NA | 2.4 | 0.81 | 74 | 2 | NA | 0.85 |

**Table 2.** List of different cathepsins which were found in the samples. The MS/MS count represents the number of MS/MS spectra leading to an identified protein. The black numbers indicate a confident identification of the protein. Proteins, which are only identified with one peptide, are not validated (number colored in grey).

| | P001 | P004 | P005 | P006 | P016 | P017 | P020 |
|---|---|---|---|---|---|---|---|
| | MS/MS count | MS/MS count | MS/MS count | MS/MS count | MS/MS count | MS/MS count | MS/MS count |
| **Cathepsin Z** | 7 | 6 | 10 | 12 | 9 | 9 | 7 |
| **Cathepsin D** | 3 | 2 | 11 | 6 | 2 | 13 | 8 |
| **Cathepsin B** | 1 | 1 | 7 | 3 | 5 | 7 | 8 |
| **Cathepsin L2** | 0 | 0 | 0 | 0 | 0 | 1 | 0 |
| **Pro-Cathepsin H** | 0 | 0 | 8 | 1 | 7 | 0 | 4 |
| **Pro-Cathepsin L** | 0 | 0 | 1 | 1 | 0 | 0 | 1 |
| **Cathepsin S** | 0 | 0 | 0 | 0 | 0 | 0 | 0 |

```
P001      GPDLTQPRSVSGSPGQSVTLSCTGTSSDVGGYNYVSWYQQHPGKAPKLMLYDVTKRPSGVPDRFSGSKSGTTASLTISGLQAEDEADYYCC
P011      EAPLTQPPSVSGAPGQRVTLSCTGSSSNLGAGWDVHWYQQLPGTVPKLLLYADRNRPSGVPERFSGSKSGTSATVAIAGLQAEDEADYYCQ
             **** ****:*** *******:**::*.   * **** **..***:**   :******:*********:*:::*:************

P001      SYAG-IDIFVLFGGGTKLTVLGQPKAAPSVTLFPPSSEELQANKATLVCLLSDFYPQVTVAWKADSSPVKAGVETTTPSKQSNNKYAASSY
P011      SYDSALSGFYVFGTGTKVIVLGQPKANPTVTIFPPSSEELQANKATLVCLLSDFYPQVTVAWKADGSPVKAGVETTKPSKQSNNKYAASSY
          ** . :. * :** ***: ******* *:**:*********************************.**********.**************

P001      LSLTPEQWKSHRSYSCQVTHEGSTVEKTVAPTECS
P011      LSLTPEQWKSHRSYSCQVTHEGSTVEKTVAPTECS
          ***********************************
```

**Figure 1.** Full length sequence alignment of the $\lambda$ light chains of our study.

```
P004a     EIVLTQSPGTLSLSPGERATLSCRASQSVSSSYLAWYQQKPGQAPRLLIYDASTRATGIPDRFSGSGSGADFLLTISSLEPEDFAMYYCQQ
P004b     EIVLSQSPDTLSLSPGERATLSCRADQSVSSNYVAWYQQKPGQAPRLLIYDAFTRATGIPDRFSGSGSGADYTLTISTLEPEDFAVYYCQQ
P006      DIQMTQSPSSLSASVGDRVTITCQASQDL-AKYLNWYQQKPGKPPKLLIYDTSNLETGVPSRFSN-GGGTDFTFTINSLQPEDLATYYCQQ
P016      DIQMTQSPSSLSASVGDRVTITCQASRDI-SNYLNWYQQKPGKAPMLLIYAASNLQTGVPSRFSGSGSGTDFTFTISSLQPEDLATYYCQQ
P017      DIQMTQSPSSLSASVGDRVTITCQASQDL-GNYLNWYQQKPGKAPRLLIYDASDLEEGVPSRFSGSGSGTDFTFTISSLQPEDFATYYCQQ
P005      DIQMTQSPSSLSASVGDRVSITCRASESI-SSYVNWYQQKPGKAPKLLIYTASSLQSGVPPRFSGSASGTDFTLTISSLQPEDFATYYCQQ
P013      DIQMTQSPSTLSASVGDAVTITCRASQSL-NVWLAWYQQKPGKPPKLLIYEASNLESGVPSRFSGSGSGTEFTLTISSLQPDDFATYYCQQ
P007      DIQMTQSPSTLSASVGDRVTITCRASQSL-SSSLAWYQQKPGKAPKLLIYDASSLETGVPSRFSGSGSGTEFTLSISSLQPDDFATYYCQH
P020      DIQMTQSPSTLSTSVGDRVTITCRASQSI-RTWLAWYQQKPGKAPKLLIYKASTLETGVPSRFSGSGSGTDFTLTISSLQPEDFATYYCQQ
          :* ::***.:** * *: .:::*:*...:    : *******: * **** :     *:* ***. .. ::: ::*.:*:*:*:* ****:

P004a     YGRS-PYTFGPGTKVDIKRTVAAPSVFIFPPSDEQLKSGTASVVCLLNNFYPREAKVQWKVDNALQSGNSQESVTEQDSKDSTYSLSSTLT
P004b     YGRS-PYTFGPGTKVDIKRTVAAPSVFIFPPSDEQLKSGTASVVCLLNNFYPREAKVQWKVDNALQSGNSQESVTEQDSKDSTYSLSSTLT
P006      YDDF-PLTFGPGTKVDIKRTVAAPSVFIFPPSDEQLKSGTASVVCLLNNFYPREAKVQWKVDNALQSGNSQESVTEQDSKDSTYSLSSTLT
P016      YGNL-PLTFGGGTKVEIKGTVAAPSVFIFPPSDEQLKSGTASVVCLLNNFYPREAKVQWKVDNALQSGNSQESVTEQDSKDSTYSLSSTLT
P017      YHTLPPLTFGGGTKVDVKRSLAAPSVFIFPPSDEQLKSGTASVVCLLNNFYPREAKVQWKVDNALQSGNSQESVTEQDSKDSTYSLSSTLT
P005      SYST-PLTFGQGTRLEIKRTVAAPSVFIFPPSDEQLKSGTASVVCLLNNFYPREAKVQWKVDNALQSGNSQESVTEQDSKDSTYSLSSTLT
P013      YNSY-PYTFGQGAKLEIKRTVAAPSVFIFPPSDEQLKSGTASVVCLLNNFYPREAKVQWKVDNALQSGNSQESVTEQDSKDSTYSLSSTLT
P007      YNSY-SLTFGQGTKVEIKRTVAAPSVFIFPPSDEQLKSGTASVVCLLNNFYPREAKVQWKVDNALQSGNSQESVTEQDSKDSTYSLSSTLT
P020      YNDY-SGTFGQGTKLEIKRTLAAPSVFIFPPSDEQLKSGTASVVCLLNNFYPREAKVQWKVDNALQSGNSQESVTEQDSKDSTYSLSSTLT
                 *** *:::::* ::**********************************************************************

P004a     LSKADYEKHKVYACEVTHQGLSSPVTKSFNRGEC
P004b     LSKADYEKHKVYACEVTHQGLSSPVTKSFNRGEC
P006      LSKADYEKHKVYACEVTHQGLSSPVTKSFNRGEC
P016      LSKADYEKHKVYACEVTHQGLSSPVTKSFNRGEC
P017      LSKADYEKHKVYACEVTHQGLSSPVTKSFNRGEC
P005      LSKADYEKHKVYACEVTHQGLSSPVTKSFNRGEC
P013      LSKADYEKHKLYACEVTHQGLSSPVTKSFNRGEC
P007      LSKADYEKHKVYACEVTHQGLSSPVTKSFNRGEC
P020      LSKADYEKHKVYACEVTHQGLSSPVTKSFNRGEC
          **********:***********************
```

**Figure 2.** Full length sequence alignment of the $\kappa$ light chains of our study.

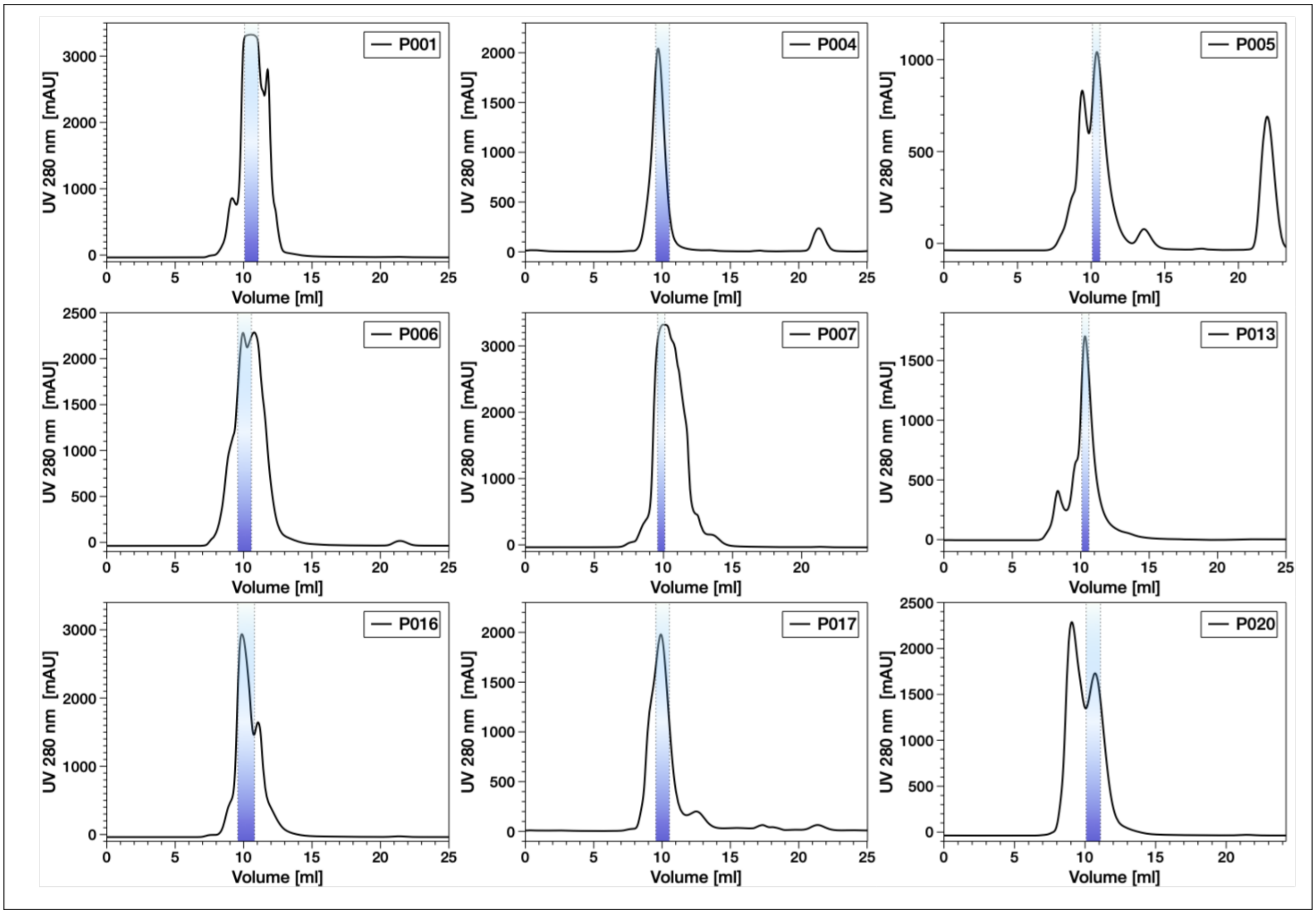


**Figure 3.** Size exclusion chromatograms of the different IgLC samples of this study. The fractions used for the remaining experiments in this study are marked in blue.

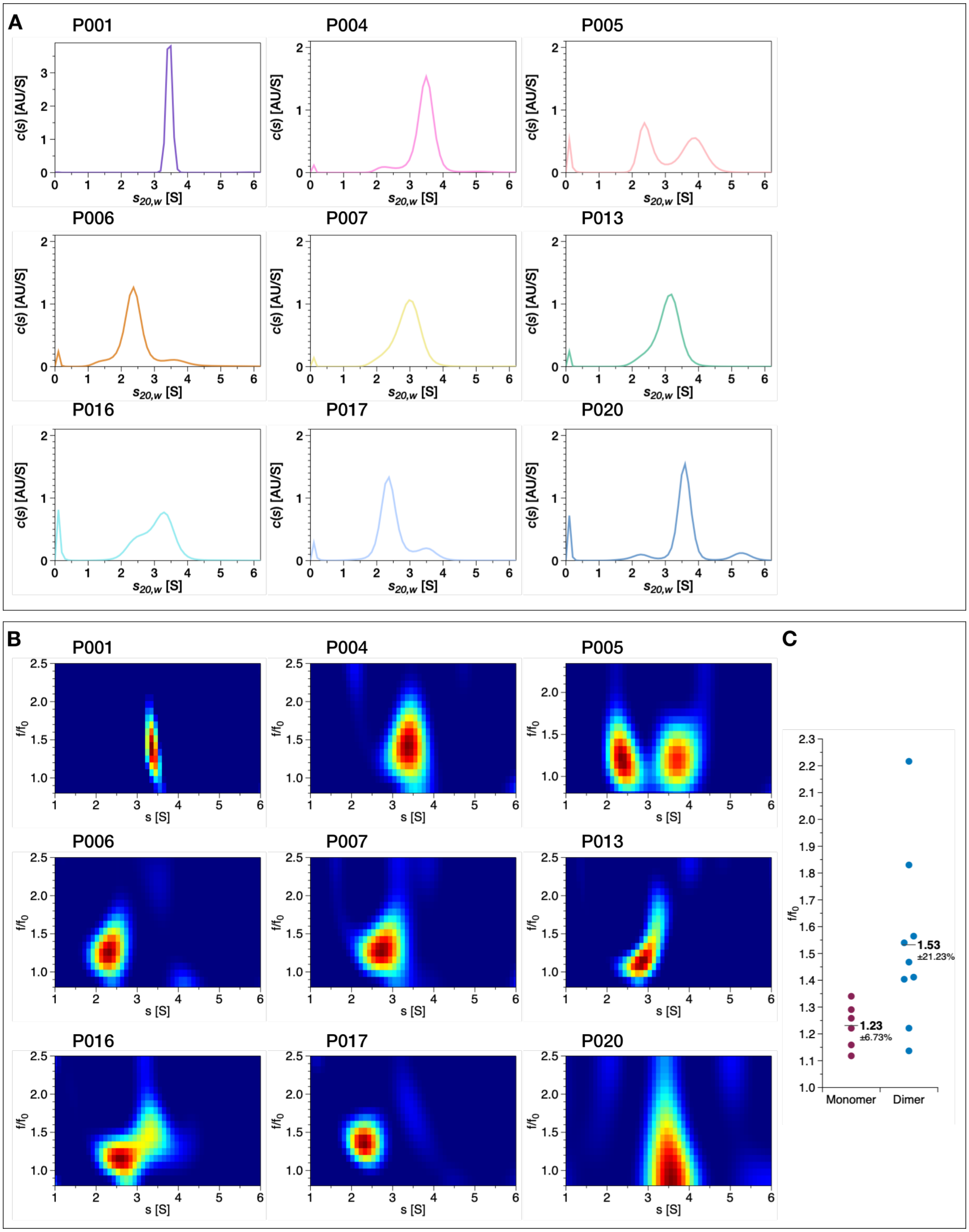


**Figure 4.** (A) Distribution of sedimentation coefficients (c($s$)) of the IgLC samples of this study and (B) application of the c($s$, $f/f0$)-model determined by sedimentation velocity experiments at 60.000 rpm. We find that the form factors of the dimers differ within the sample set. The dimers can appear very globular e.g. P020 with a $f/f_0$ of 1.14 or elongated such as P006 $f/f_0$ of 2.22 and P007, P013 and P017 $f/f_0$ between 1.54 and 1.83. However the signal for the dimer of P006 is very low. (C) shows the variation of $f/f_0$ between the dimers compared to the monomers. The monomer of P004 and P020 was excluded due to the low signal.

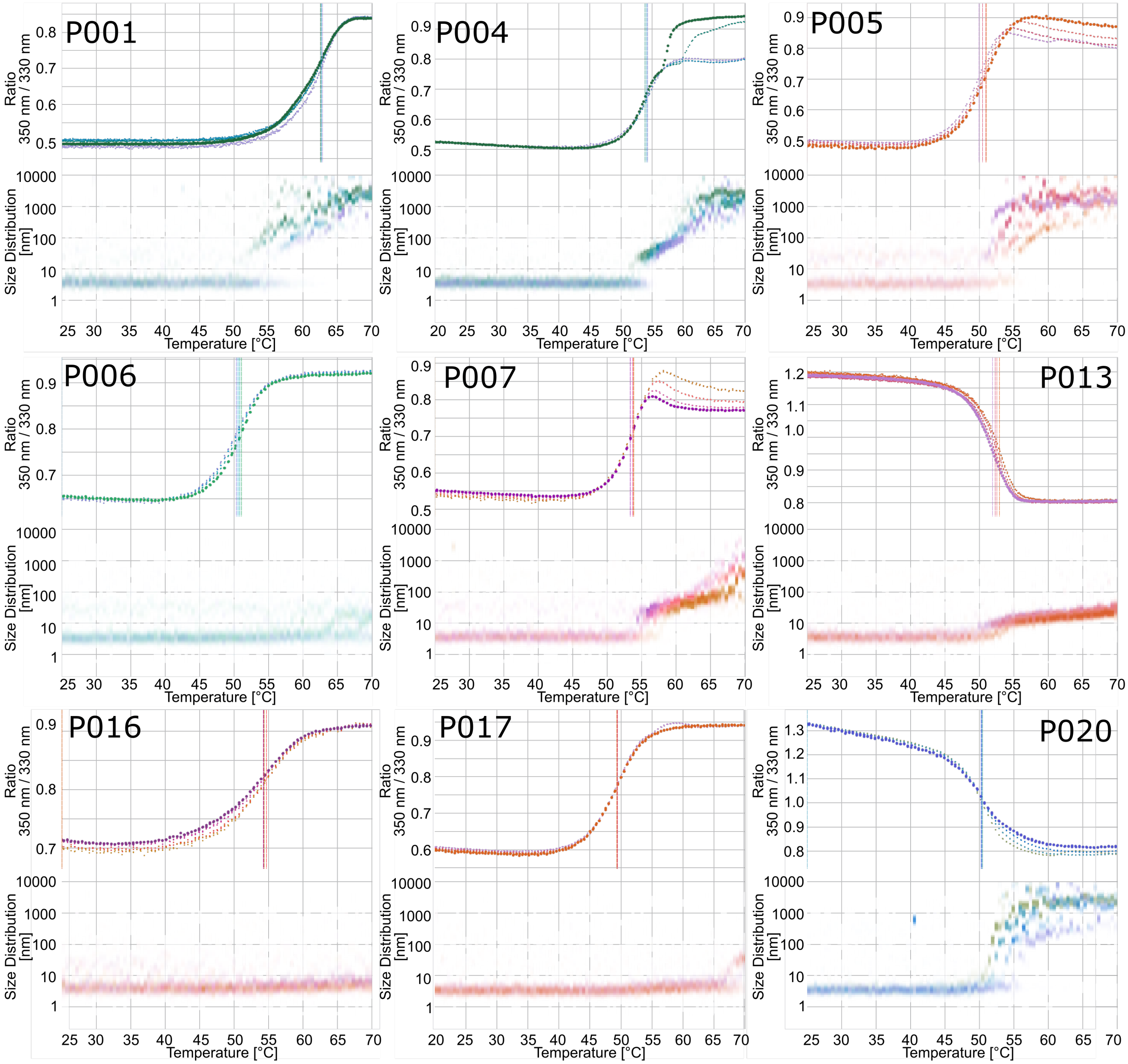


**Figure 5.** Concentration dependent thermal unfolding experiments of the IgLC samples of our study. The samples were scanned from 25-70°C at 1°C per minute. In each case, the evolution of the ratio of the intrinsic fluorescence intensities at 350 and 330 nm is shown on top, and the evolution of the size distribution, measured by dynamic light scattering (DLS) is shown on a logarithmic scale on the bottom. Each sample was measured at 4 concentrations, as the undiluted stock solution as well as 3 dilutions by a factor of 2 each. The concentrations of the stock solutions are 97 $\mu$M (P001), 150 $\mu$M (P004), 91 $\mu$M (P005), 139 $\mu$M (P006), 103 $\mu$M (P007), 97 $\mu$M (P013), 99 $\mu$M (P016), 158 $\mu$M (P017) and 81 $\mu$M (P020)

.

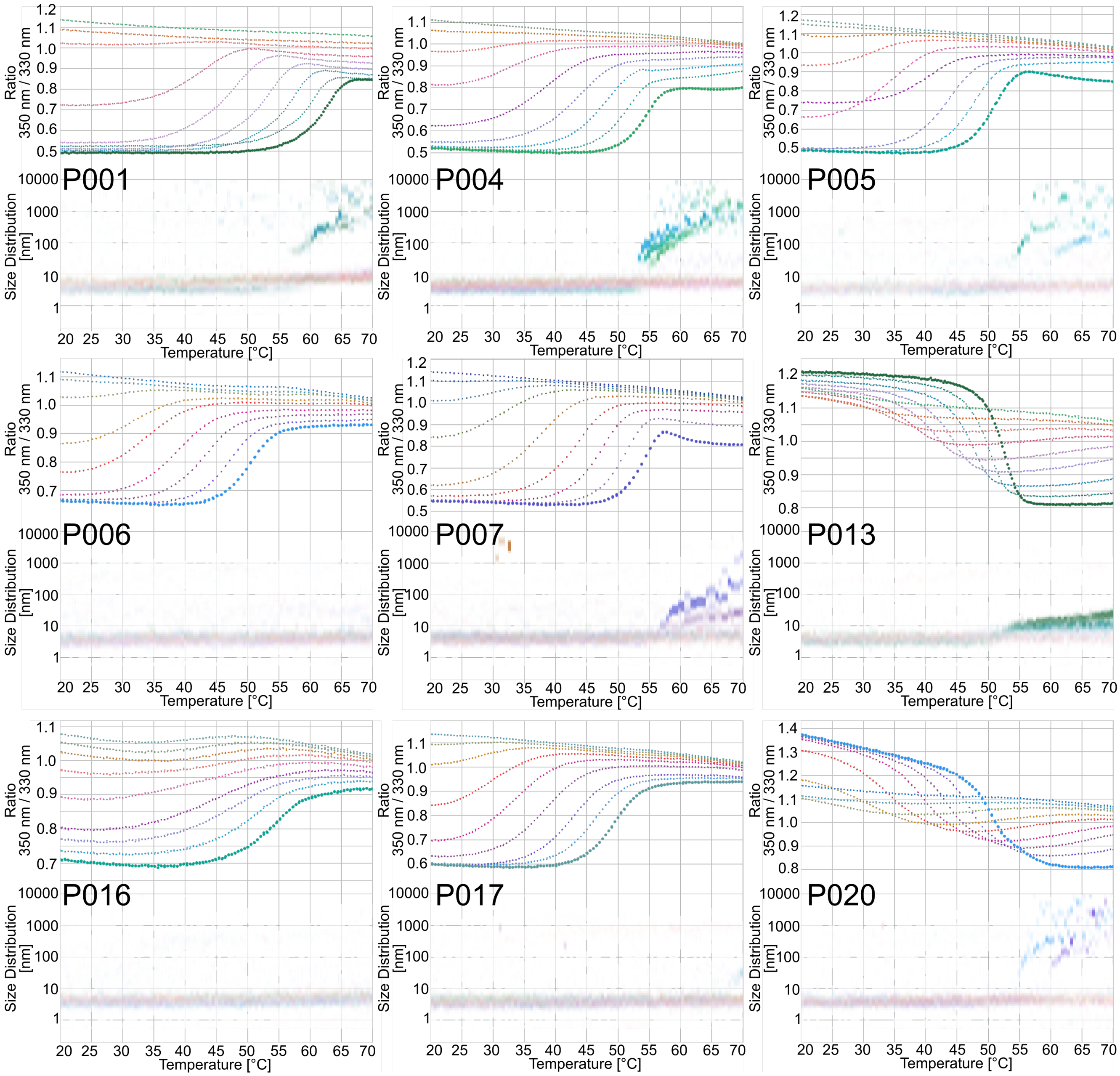


**Figure 6.** Temperature-dependent chemical denaturation of the IgLC samples of this study by urea. In each case, the evolution of the ratio of the intrinsic fluorescence intensities at 350 and 330 nm is shown on top, and the evolution of the size distribution, measured by DLS is shown on a logarithmic scale on the bottom. The protein concentration corresponds to a 5-fold dilution of the stock solution used for the thermal denaturation in Figure 5. The urea concentrations are in each case 0, 0.67,1.34, 2.01, 2.68, 3.35, 4.02, 4.69, 5.36 M. The samples were scanned from 20-70°C at 1°C per minute.

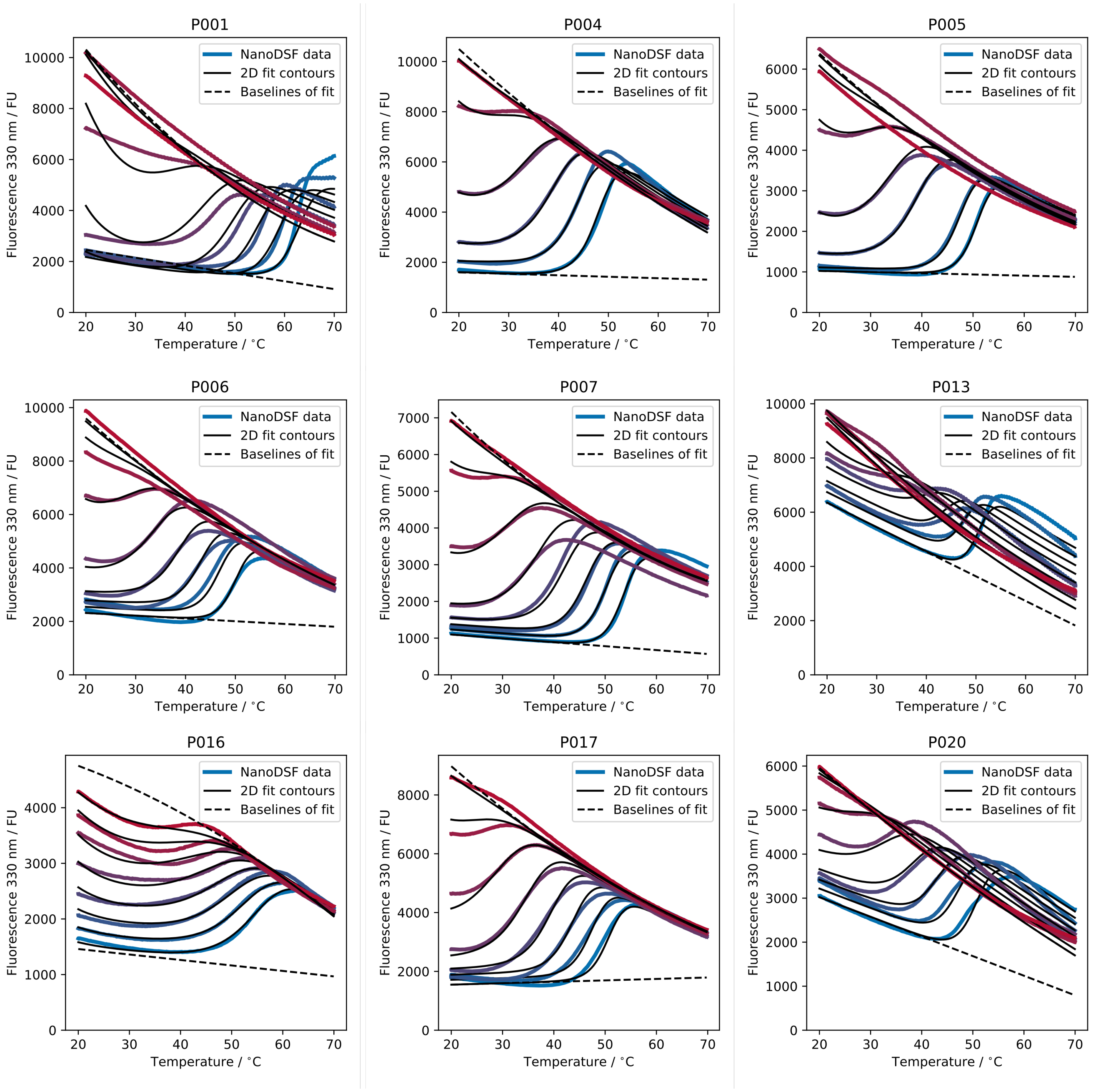


**Figure 7.** Global fits of the temperature-dependent chemical denaturation of the IgLC samples of this study by urea. The data is the same as in the previous figure, but instead of using the fluorescence intensity ratio, the fits are performed simultaneously on the fluorescence intensities at both 330 nm (shown here) and 350 nm.

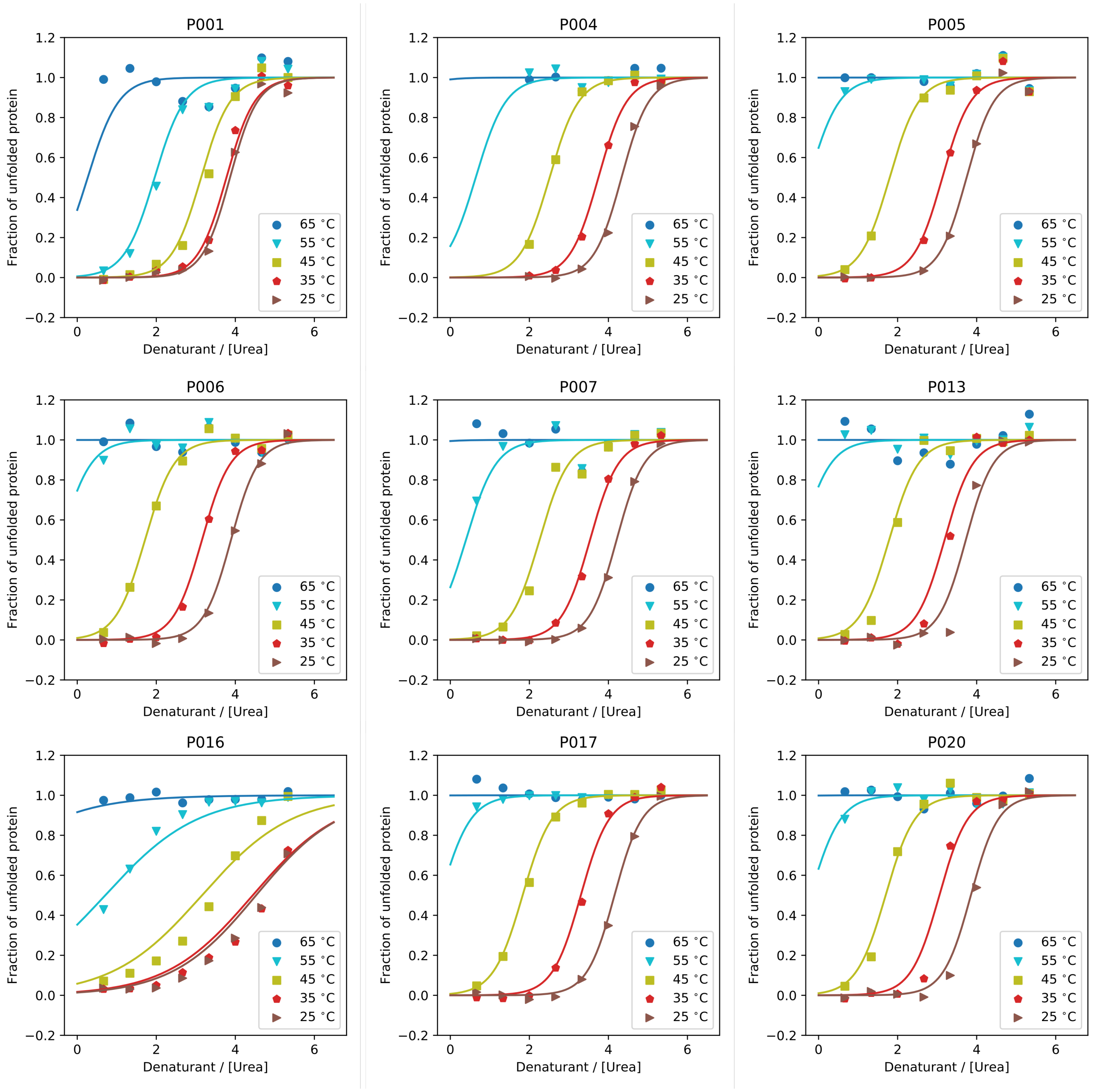


**Figure 8.** Results from the global fits of the temperature-dependent chemical denaturation of the IgLC samples of this study by urea. Shown is the fraction of unfolded protein as a function of urea concentration at different temperatures.

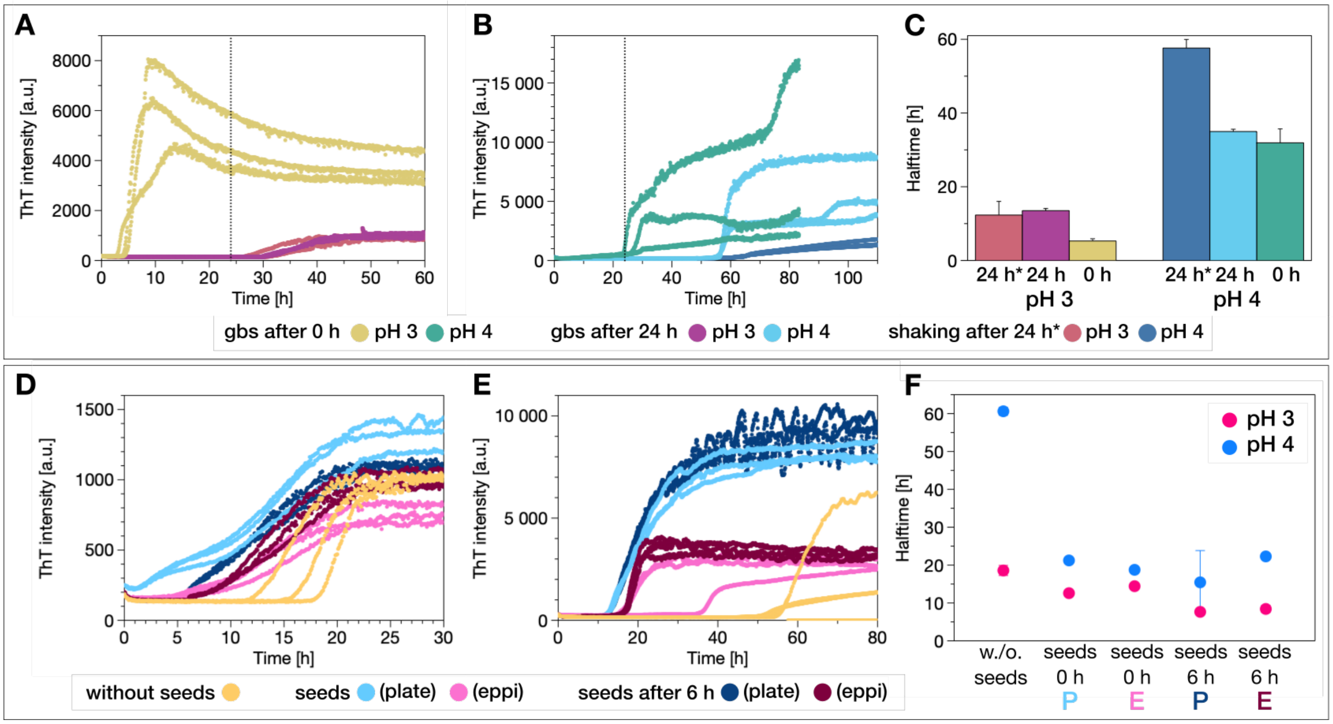


**Figure 9.** Aggregation assays of P016 at (A) pH 3 and (B) pH 4 monitored in a high-binding surface plate in the presence of glass beads and conditions of mechanical agitation and with addition of glass beads after 24 h pre-incubation without shaking and the (C) aggregation halftimes (top). Aggregation assays of P016 at (D) pH 3 and (E) pH 4 in a high-binding plate under quiescent conditions. Seeds prepared in a high-binding plate and prepared in an Eppendorf tube are added at the beginning and after 6 h pre-incubation and (F) the halftimes are analysed. The pre-incubation times (24 or 6 h) are subtracted from the halftimes.

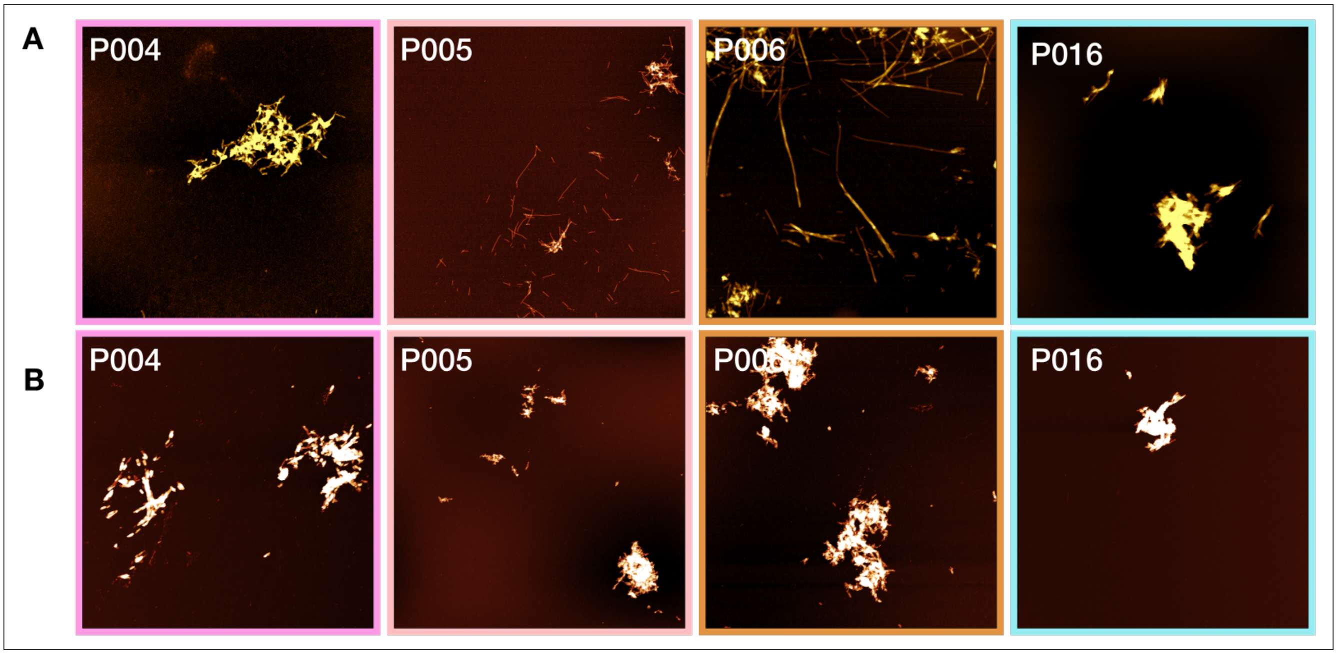


**Figure 10.** AFM-height-images of aggregates prepared in different reaction vessels. We used as seeds both fibrils which had been prepared in a high-binding surface plate in the presence of glass beads (A), as well as fibrils which had been prepared in the same volume in an 2 ml Eppendorf tube with glass beads (B). The presence of fibrils was confirmed using atomic force microscopy. Although the presence of fibrils could be confirmed in both setups, the total ThT-fluorescence intensity was lower if seeds prepared in an Eppendorf tube were used. The image scale is 5 x 5 $\mu$m. The colour range represents the height from -2 to 15 nm.

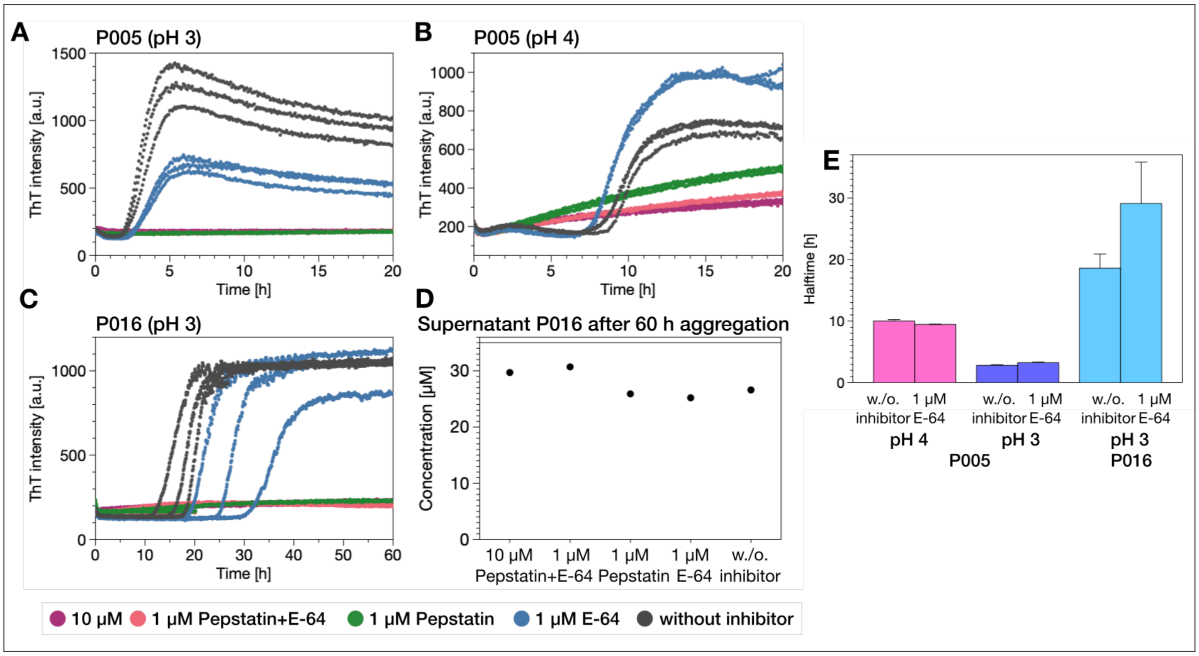


**Figure 11.** Influence of two different protease inhibitors pepstatin A and E-64 on the aggregation kinetics of the samples P005 at (A) pH 3 and (B) pH 4 and (C) P016 at pH 3 monitored in a high-binding plate under agitation conditions. (D) AFM-height-images of aggregated P016 at pH 2 (purple), pH 4 (blue), pH 3 (cyan) and pH 3 in presence of 10 $\mu$M pepstatin A and E-64 after 70 h and the extracted height profiles of different fibrils (top left) and the width of the twist (bottom left). Fibrils at pH 3 do not display any twist. The image scale is 5 x 5 $\mu$m. The colour range represents the height from -2 to 15 nm.

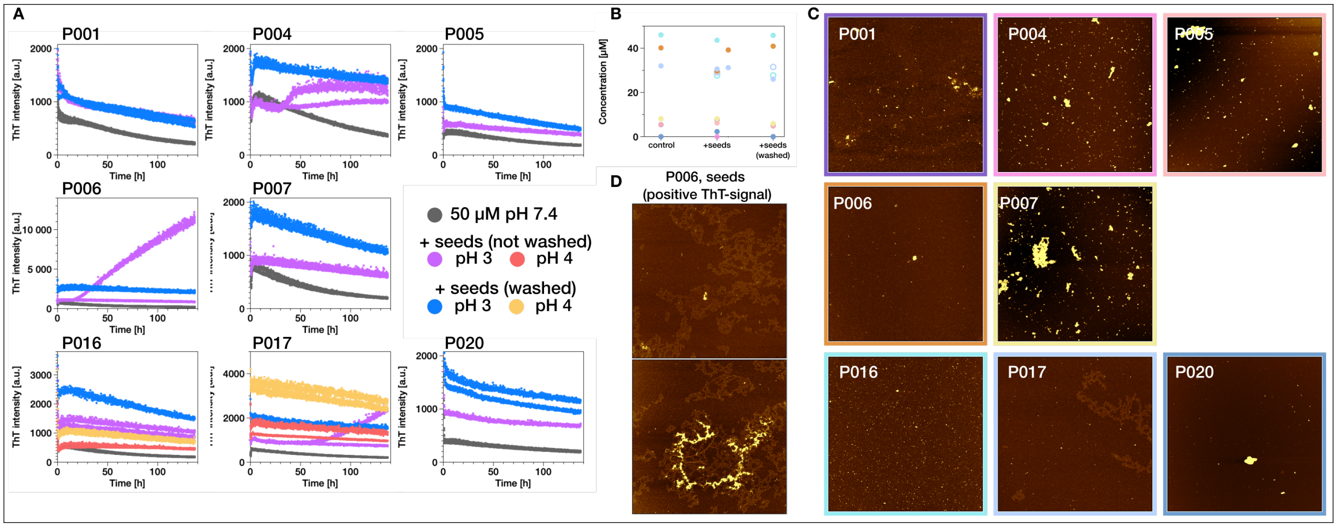

**Figure 12.** (A) Aggregation experiment at 55°C monitored in a non-binding surface plate under agitation conditions in the presence of glass beads. 5% Seeds which were produced at pH 3 or pH 4 in an Eppendorf tube were added to 50 $\mu$M light chain. (B) The soluble content was determined after the experiment by UV-absorbance. AFM-height-images of the light chains at the end of the experiments (C) and of aggregated P006 with the positive ThT-signal (D). Fibrils at pH 3 do not display any twist. The image scale is 5 x 5 $\mu$m. The colour range represents the height from -2 to 5 nm.

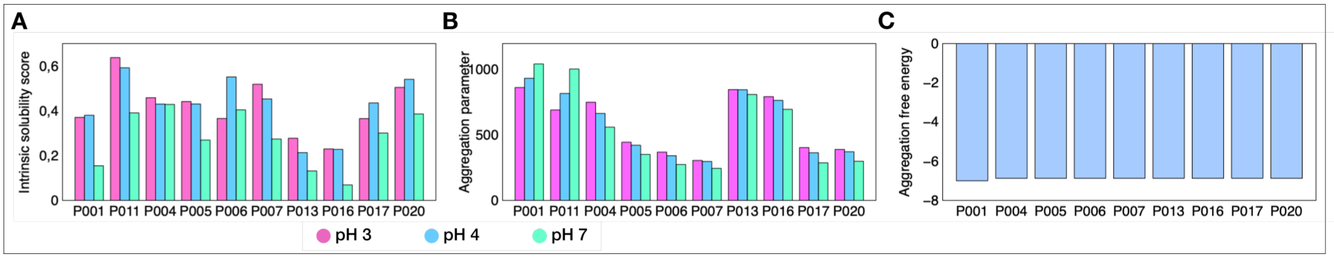

**Figure 13.** (A) Intrinsic solubility score measured by CamSol and (B) aggregation parameter determined by the Tango algorithm at pH 3, pH 4 and pH 7, (C) aggregation free energy computed with the Pasta algorithm.

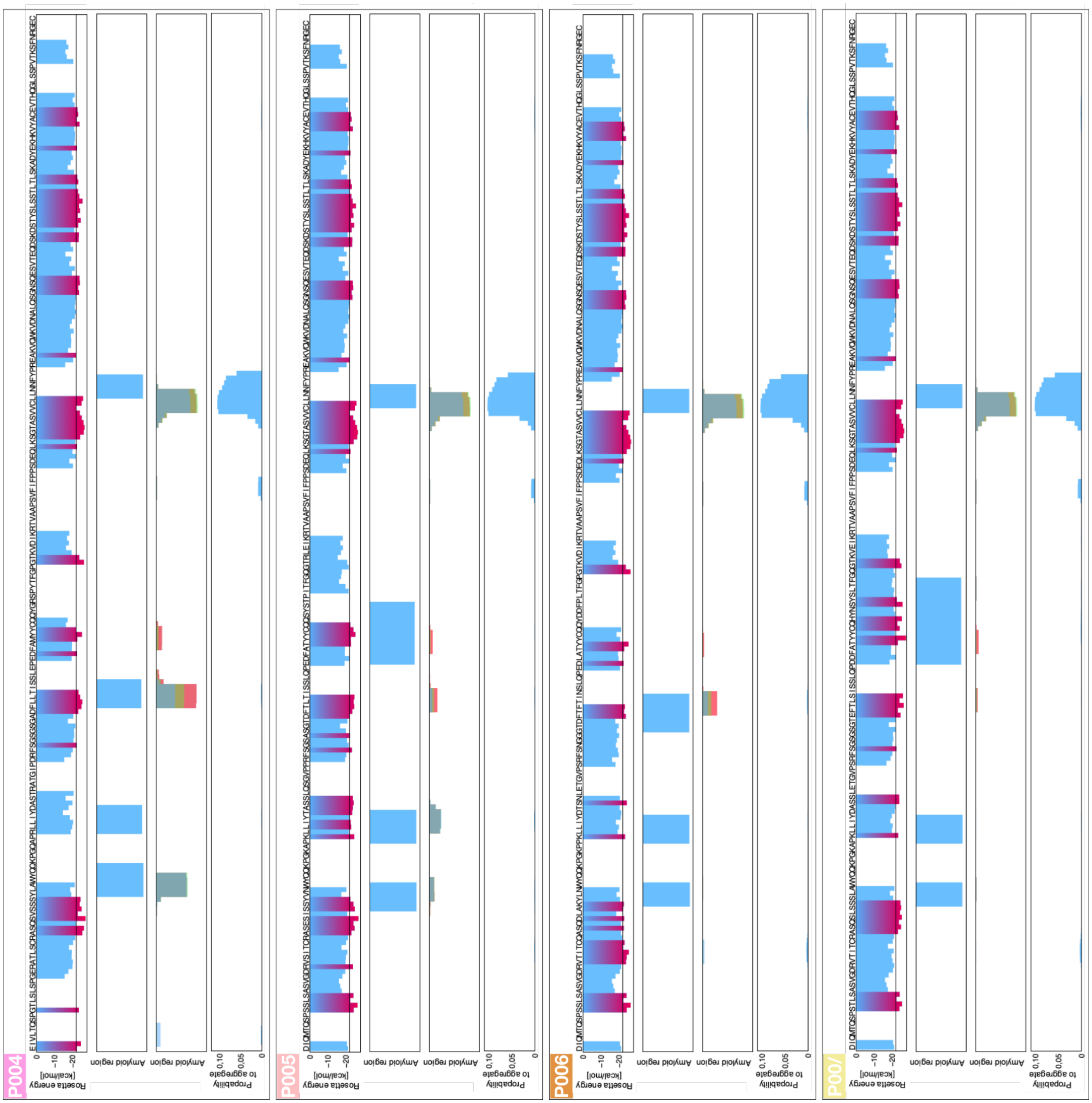


**Figure 14.** The $\kappa$ IgLC sequences P004, P005, P006 and P007 were analysed with different amyloid prediction tools and the results visualised on the sequence. Rosetta energy was determined by ZipperDB, amyloidogenic sequence regions with the algorithms Waltz, Tango and Pasta. The analyses with Tango were performed at pH 3 (red), pH 4 (green) and pH 7 (blue).

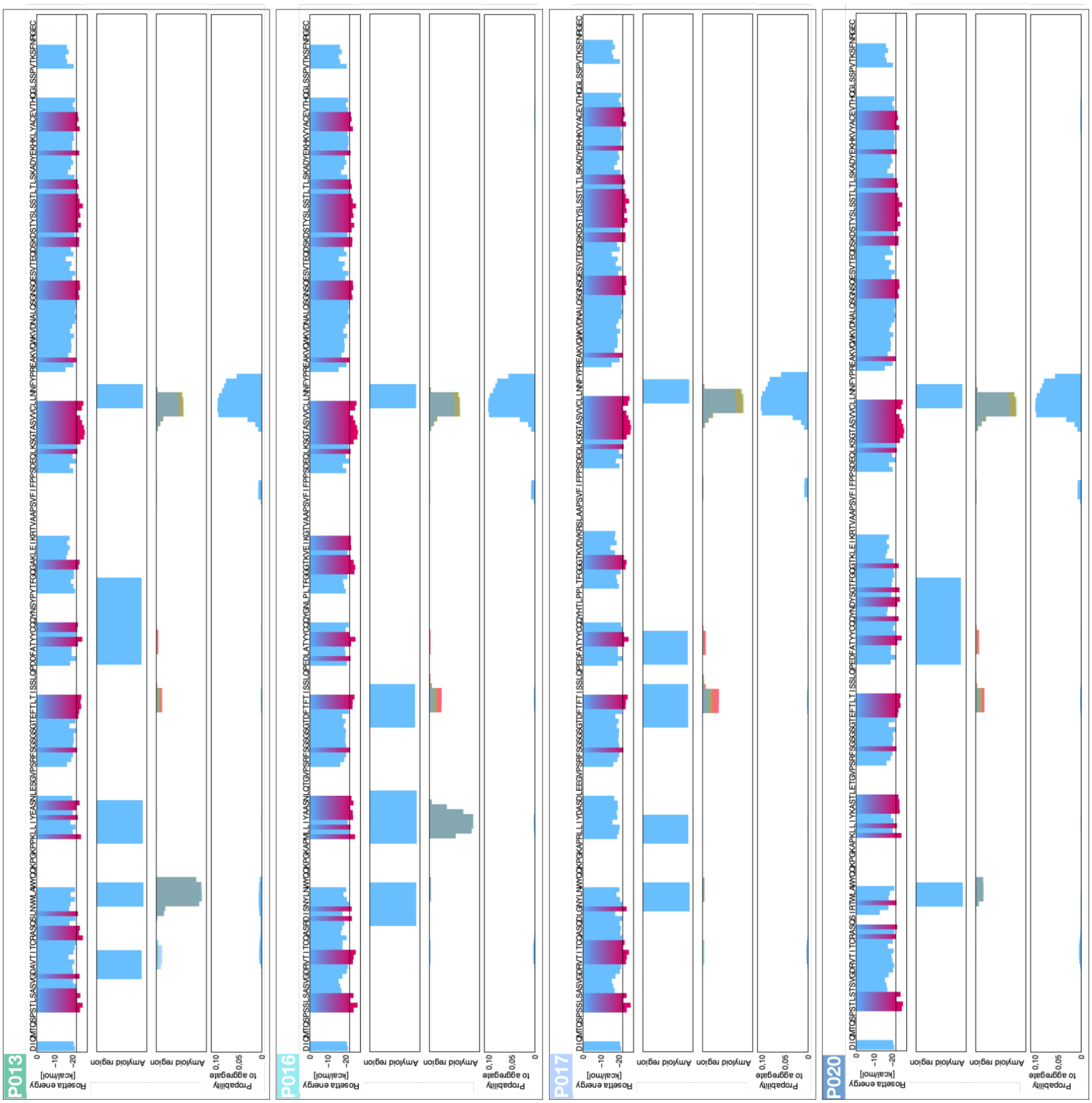


**Figure 15.** The $\kappa$ IgLC sequences P013, P016, P017 and P020 were analysed with different amyloid prediction tools and visualised on the sequence. Rosetta energy was determined by ZipperDB, amyloidogenic sequence regions with the algorithms Waltz, Tango and Pasta. The analyses with Tango were performed at pH 3 (red), pH 4 (green) and pH 7 (blue).

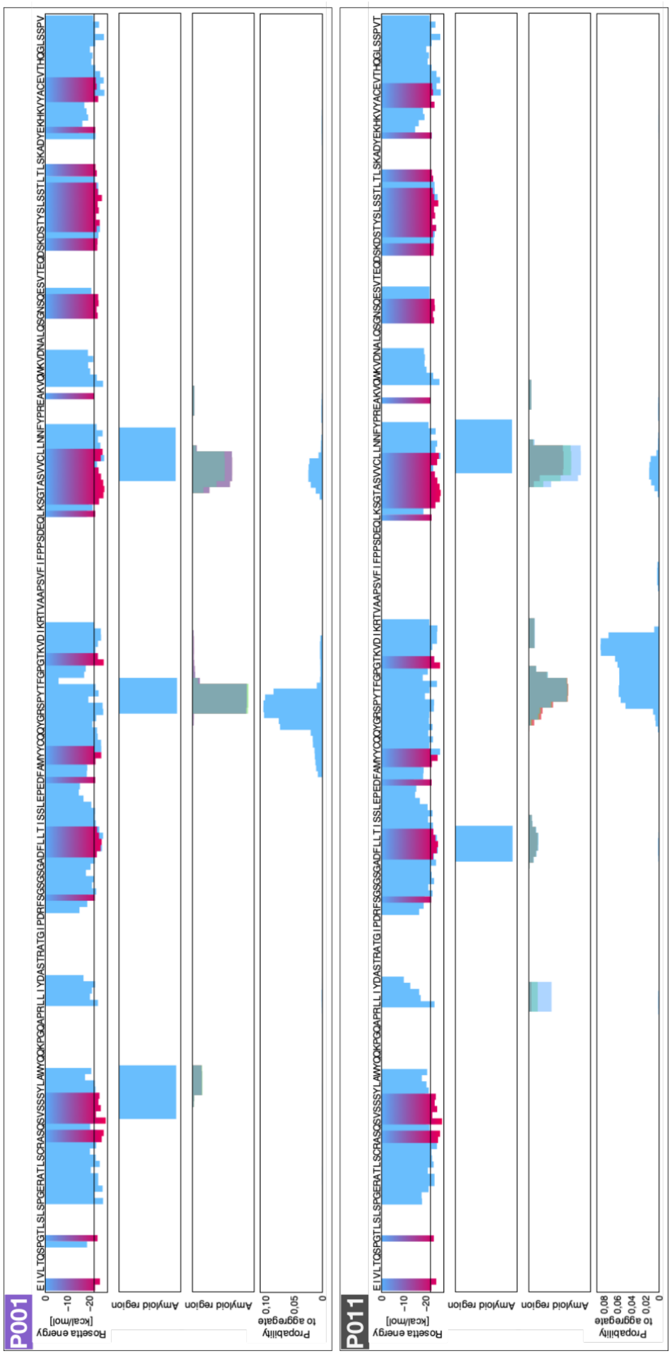


**Figure 16.** The $\lambda$ IgLC sequences P001 and P011 were analysed with different amyloid prediction tools and visualised on the sequence. Rosetta energy was determined by ZipperDB, amyloidogenic sequence regions with the algorithms Waltz, Tango and Pasta. The analyses with Tango were performed at pH 3 (red), pH 4 (green) and pH 7 (blue).

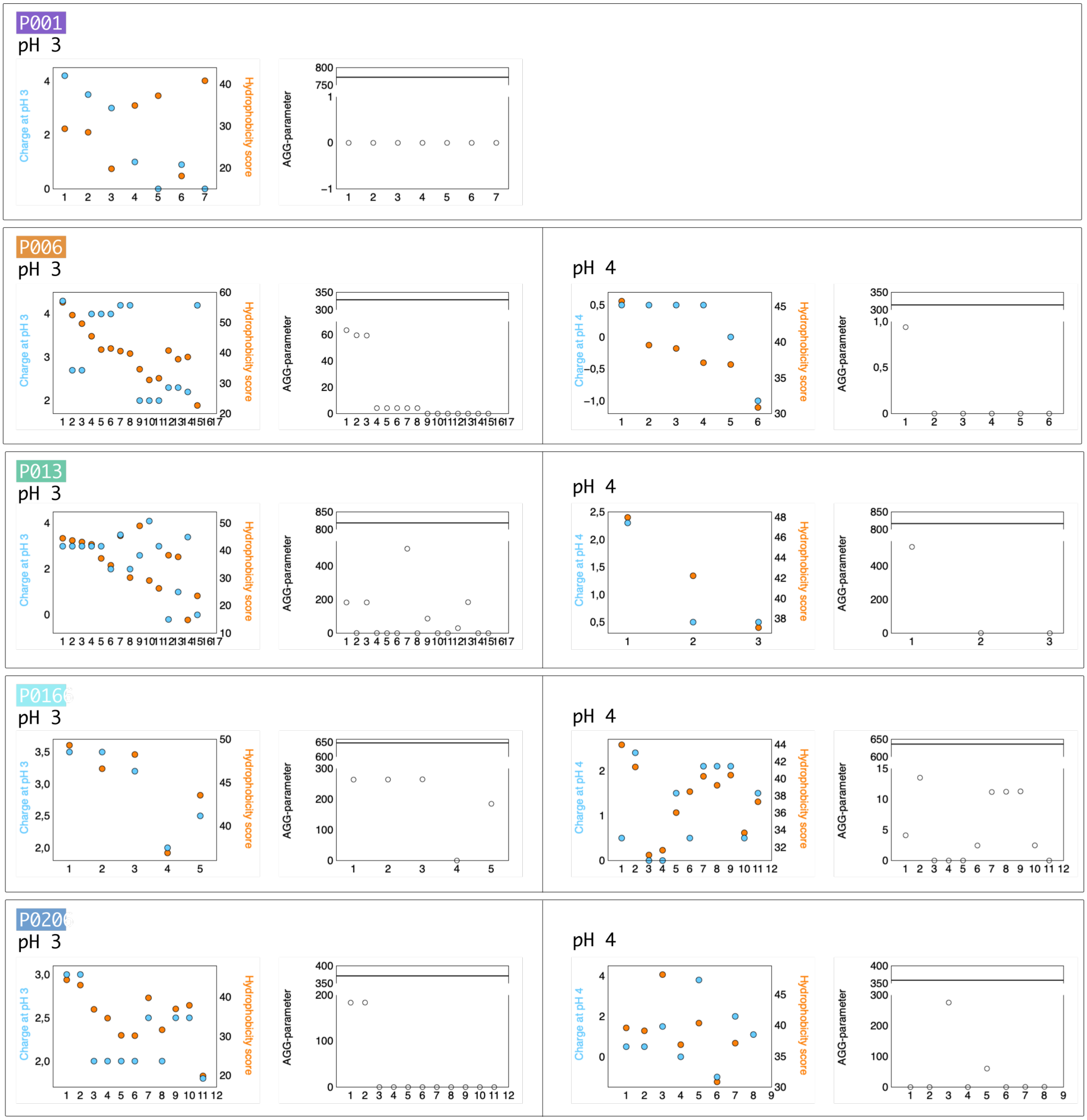


**Figure 17.** The sequence regions of P001, P006, P013, P016 and P020 that were found in the aggregates formed at acidic pH values and identified by mass spectrometry. A complete list of all fragments and their masses is provided as a separate pdf file attached to this submission. The aggregation products formed in Eppendorf tubes under agitation conditions were centrifuged using an Optima MAX-XP ultracentrifuge (Beckman Coulter) in a TLA-55 rotor at 40,000 rpm at 20° for 45 min. The pellet was re-suspended in 150 mM citric acid (pH 3 or pH 4) and centrifuged again for 45 min. This washing procedure to remove the soluble fragments was conducted three times. The washed aggregates were dissolved in 6 M urea and subsequently measured on a . The sequence regions were analysed according to their charge at the depicted pH value (blue) and hydrophobicity score (orange) using the Peptide Analyzing Tool from Thermofisher. The hydrophobicity score is based on the index proposed by Krokhin and Spicer[1]. The AGG parameters of the sequence fragments were analysed using the TANGO algorithm and the AGG parameter of the full length seqeunce is indicated with a horizontal line.

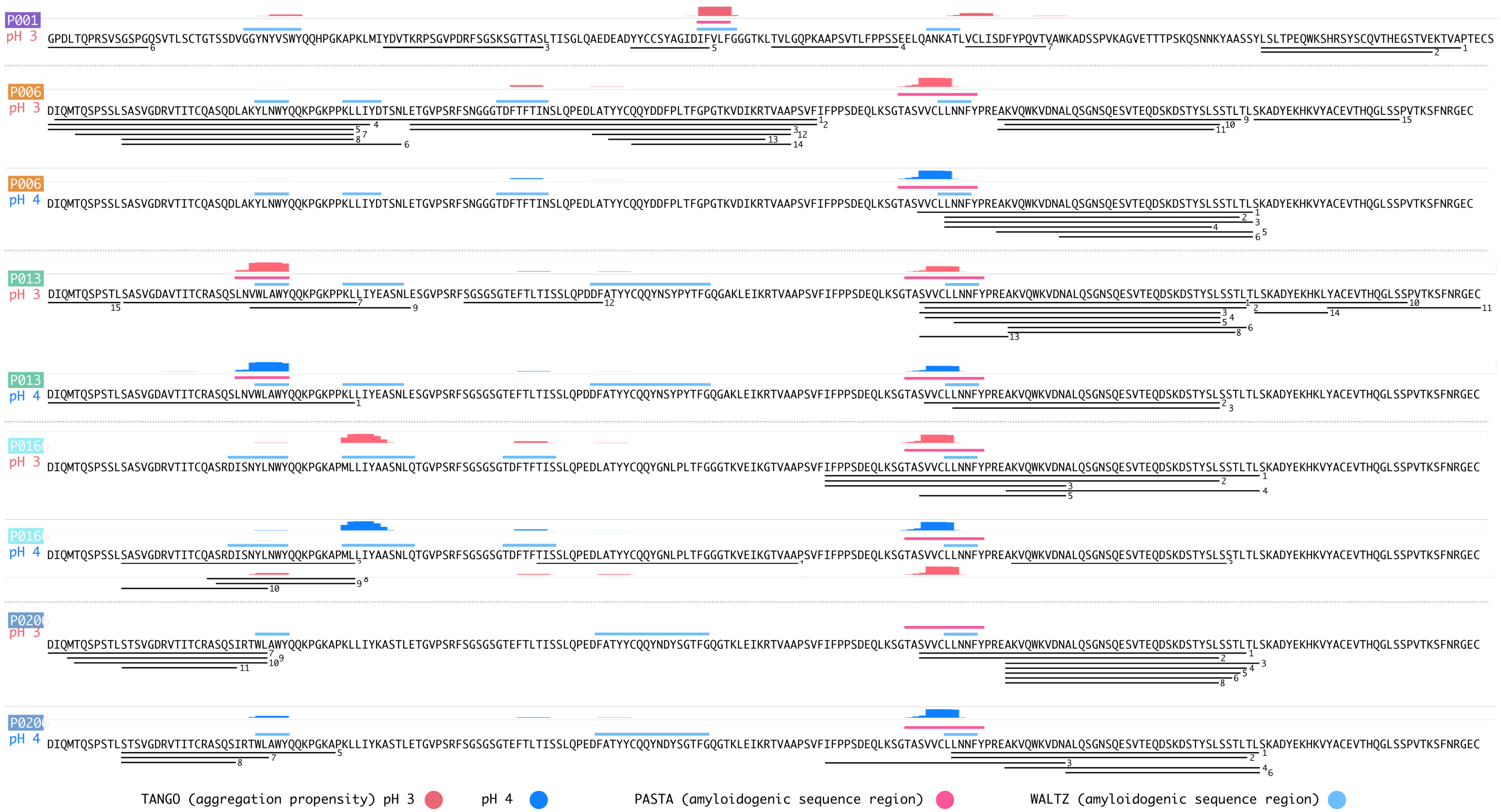


**Figure 18.** The IgLC sequences P001, P006, P013, P016 and P020 were analysed with different amyloid prediction tools and the predicted aggregation hot spots visualised on the sequence. Amyloid propensity was determined with the algorithm PASTA (red pH 3, blue pH 4) and the amyloidogenic sequence regions with the algorithms TANGO and WALTZ. The sequence parts which were found in the aggregates formed at acidic pH values are marked with black lines (figure 17.

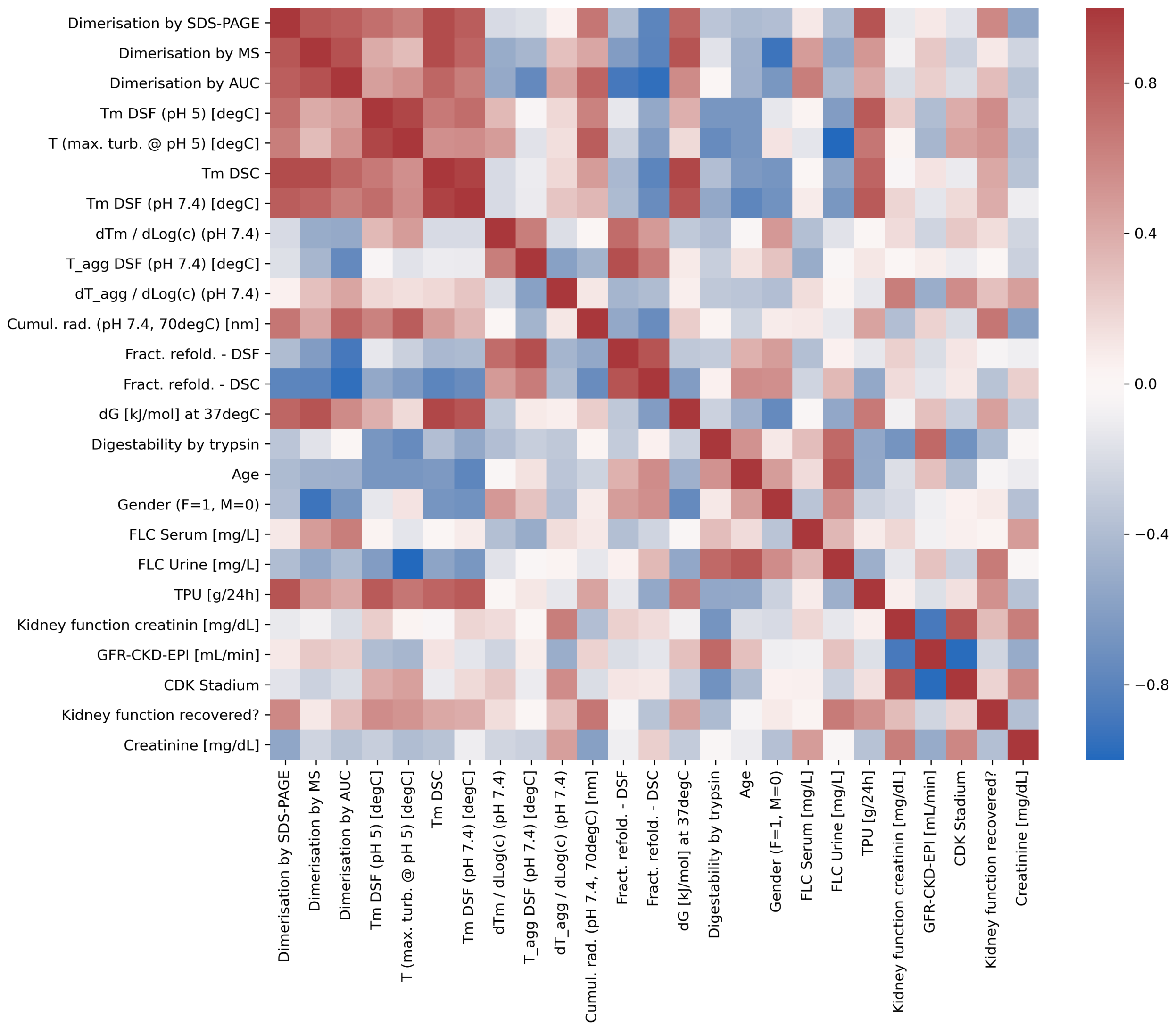


**Figure 19.** Pearson correlation matrix between all experimental parameters of this study and clinical patient descriptors.

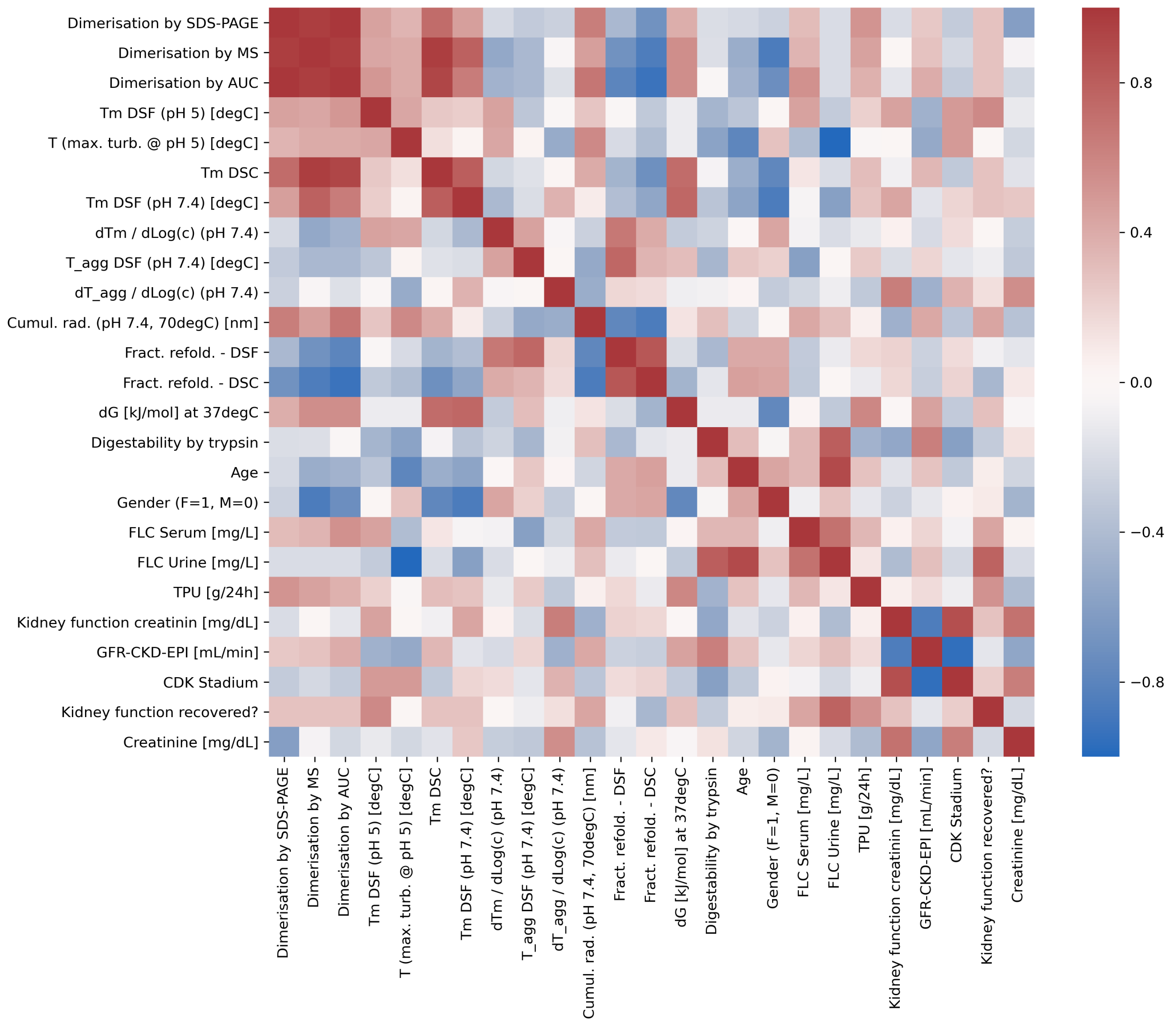


**Figure 20.** Spearman correlation matrix between all experimental parameters of this study and clinical patient descriptors.

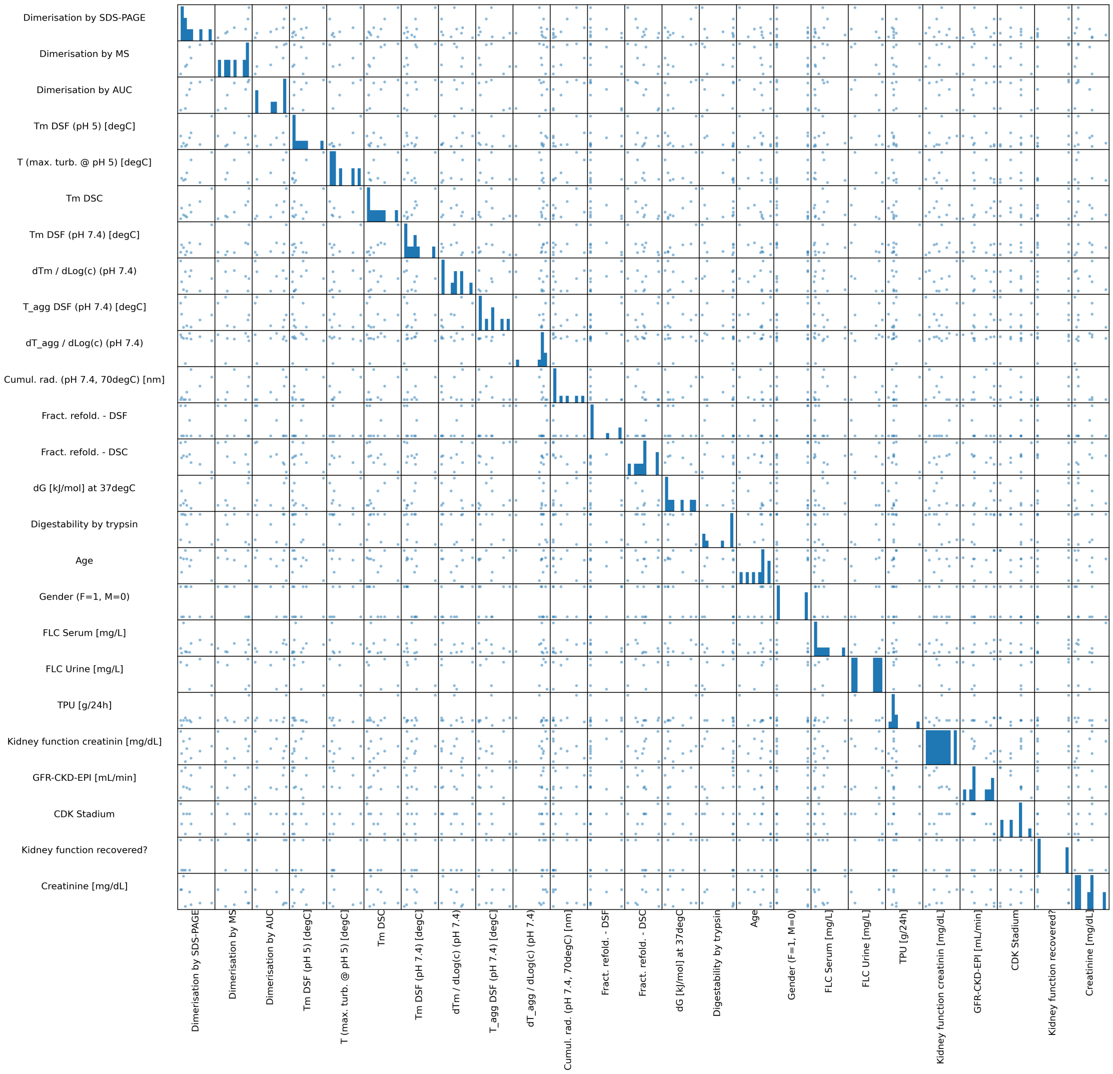


**Figure 21.** Scatter matrix that shows the pairwise correlation plots in the off-diagonal elements, and a histogram of the parameter values in the diagonal elements.

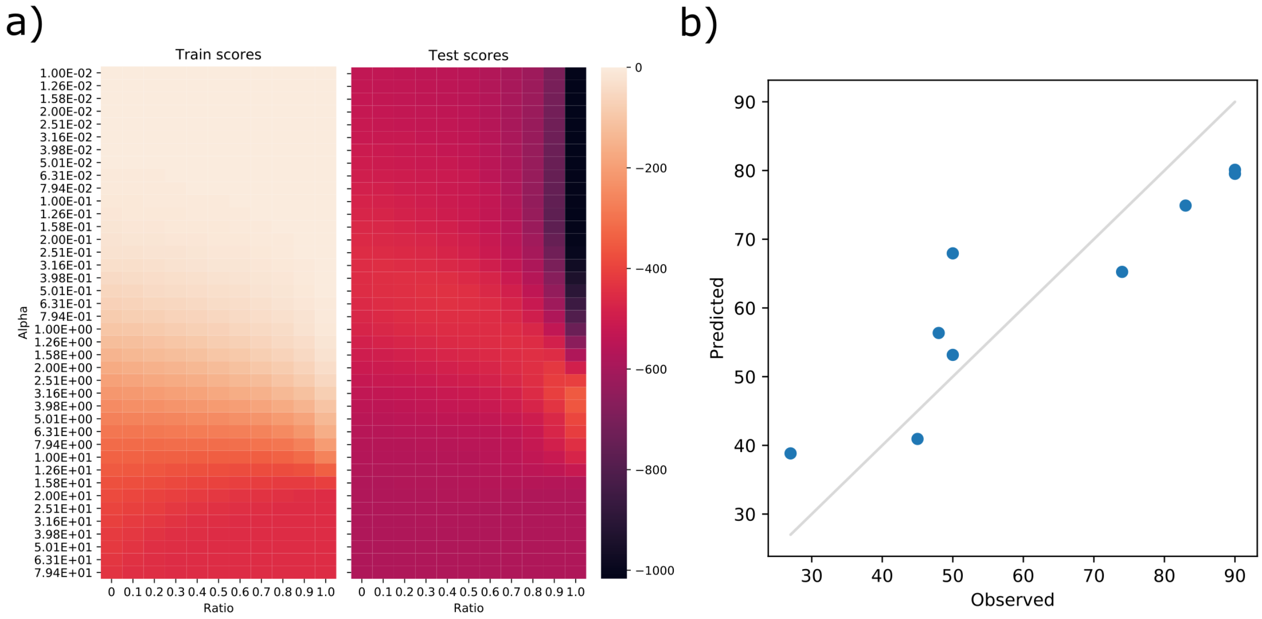


**Figure 22.** An elastic net model was trained to predict (GFR-CKD-EPI) using all the biophysical parameters measured of the light chains. a) Negative mean squared errors of the models are shown for training and test sets using 4-fold cross validation as a function of the regularisation parameters. No data points were omitted as a separate validation set because of the low amount of data. b) The best model with $Alpha = 3.16$ and $L1Ratio = 1.0$ is shown. This model uses only the three parameters trypsin digestibility, ΔG, and $\mathrm{d}T_{agg}/\mathrm{dlog}\,c$, with a Pearson correlation of 0.91. The mean error of cross validation of the model is 18.76±15.1. The influence of $\mathrm{d}T_{agg}/\mathrm{dlog}\,c$ seems to mainly arise from a single data point, so the two other parameters are probably more robust. This type of behaviour emphasises the need for significantly larger data sets than the ones that are currently available. Searching for correlations within combinations of parameters provides a large freedom that needs to be constrained by the size of the data set.

#### References

**1.** Krokhin, O. V. & Spicer, V. Peptide retention standards and hydrophobicity indexes in reversed-phase high-performance liquid chromatography of peptides. *Anal. chemistry* **81**, 9522–9530 (2009).
